# Logic of brain-wide connections with the cerebellar cortex

**DOI:** 10.64898/2026.05.29.728551

**Authors:** Sam Clothier, Dimitar Kostadinov, Beverley A. Clark, Michael Häusser, L. Federico Rossi

## Abstract

The cerebellum governs movement and cognition through multiple lobules that carry distinct signals and connect to distinct long-range brain networks. How these specialized computations are reconciled with the brain-wide convergence required for coherent behaviour is unknown, in part because the input-output connectivity of functionally distinct lobules has not been mapped. We combined anterograde and retrograde transsynaptic viral tracing with whole-brain two-photon tomography to map the monosynaptic and disynaptic inputs and outputs of Lobule V (LV) and Simplex (LS), two lobules preferentially involved in movement and reward, respectively. Monosynaptic connectivity was strongly segregated: LV and LS received climbing fiber and mossy fiber inputs from largely non-overlapping source regions and projected to distinct targets within the cerebellar nuclei. In contrast, disynaptic inputs to LV- and LS-projecting olivary neurons arose from a shared set of forebrain, midbrain, and hindbrain regions, with the segregation re-emerging at the level of fine-scale spatial topography within these regions. Disynaptic outputs through the cerebellar nuclei likewise overlapped more than the monosynaptic projections to the nuclei themselves. These findings reveal that overlapping brain-wide pathways interface with locally specialized monosynaptic cerebellar pathways. This organizational logic could support distinct movement- and reward-related computations in the cerebellum while distributing outputs to shared downstream networks, providing an architectural substrate for both segregating and integrating cerebellar motor and cognitive functions.

## Introduction

The canonical circuit organization of the cerebellum has led to the prevailing idea that it performs a defined transformation (Eccles et al., 1967): contextual information provided by mossy fiber (MF) input is conditioned by instructive signals provided by climbing fibers (CFs) and funneled to the rest of the brain via convergent Purkinje cell output to cerebellar nuclear neurons (Apps & Garwicz, 2005; Herzfeld & Lisberger, 2025; Raymond & Medina, 2018). This local circuit motif is repeated as ‘modules’ that tile the cerebellum and represent its fundamental processing units (Apps et al., 2018).

The homogeneous local organization of cerebellar modules contrasts with the heterogeneity of cerebellar input and output pathways linking these modules to the rest of the brain (Fujita et al., 2020; Kelly & Strick, 2003; Pisano et al., 2021). These diverse long-range connections, which link the cerebellum to both motor and non-motor brain regions, suggest that cerebellar computations can be applied across a broad range of behavioral contexts through connections between specific cerebellar regions and specific long-range partners (Ito, 2008; Kelly & Strick, 2003; Kostadinov & Häusser, 2022; Li & Mrsic-Flogel, 2020; Zhu et al., 2023).

Despite the crucial role that long-range connections play in orchestrating cerebellar modules, there is no cerebellar cortical region for which we have a comprehensive map of the relationship between local information processing and MF input pathways, CF input pathways, and target regions in the cerebellar nuclei (CbN). While functional mapping is revealing an increasingly detailed organisation of the cerebellar cortex in modules and microzones (Apps & Hawkes, 2009; Kostadinov et al., 2019), only recent optogenetic (Proville et al., 2014) and transynaptic viral tracing (Huang et al., 2013; Pisano et al., 2021; Wang et al., 2022) tools have enabled the mapping of polysynaptic pathways with sufficient anatomical granularity(Apps & Hawkes, 2009). Thus, it remains unclear how the diversity of functional properties among cerebellar modules is related to their long-range connectivity.

To address this knowledge gap, we have created a comprehensive map of the input pathways and output targets of two cerebellar cortex subregions involved in governing forelimb movements: Lobule V (LV) and Simplex (LS) (Heffley et al., 2018; Kostadinov et al., 2019; Kostadinov et al., 2025; Lee et al., 2015; Wagner et al., 2021). We focused on these regions because, while they share some similarities, they exhibit clear functional biases: instructive signals in LV preferentially encode movement, whereas instructive signals in LS preferentially encode reward (Kostadinov et al., 2025).

We reasoned that these lobules could use different connectivity logic to implement these computations. At one extreme, they may participate in completely discrete or segregated pathways: receiving information from distinct sets of areas and feeding their computations into distinct downstream networks (**Figure 1a,b**). At the other extreme, these regions may participate in integrated pathways: receiving shared inputs from an overlapping set of areas and feeding their computations to overlapping downstream networks that may utilize both reward and motor error signals (**Figure 1c,d**) (Finkelstein et al., 2026). These hypotheses sit at opposing ends of a continuum, and the actual organisation likely combines features of both - the question is at what spatial scale.

**Figure 1:**
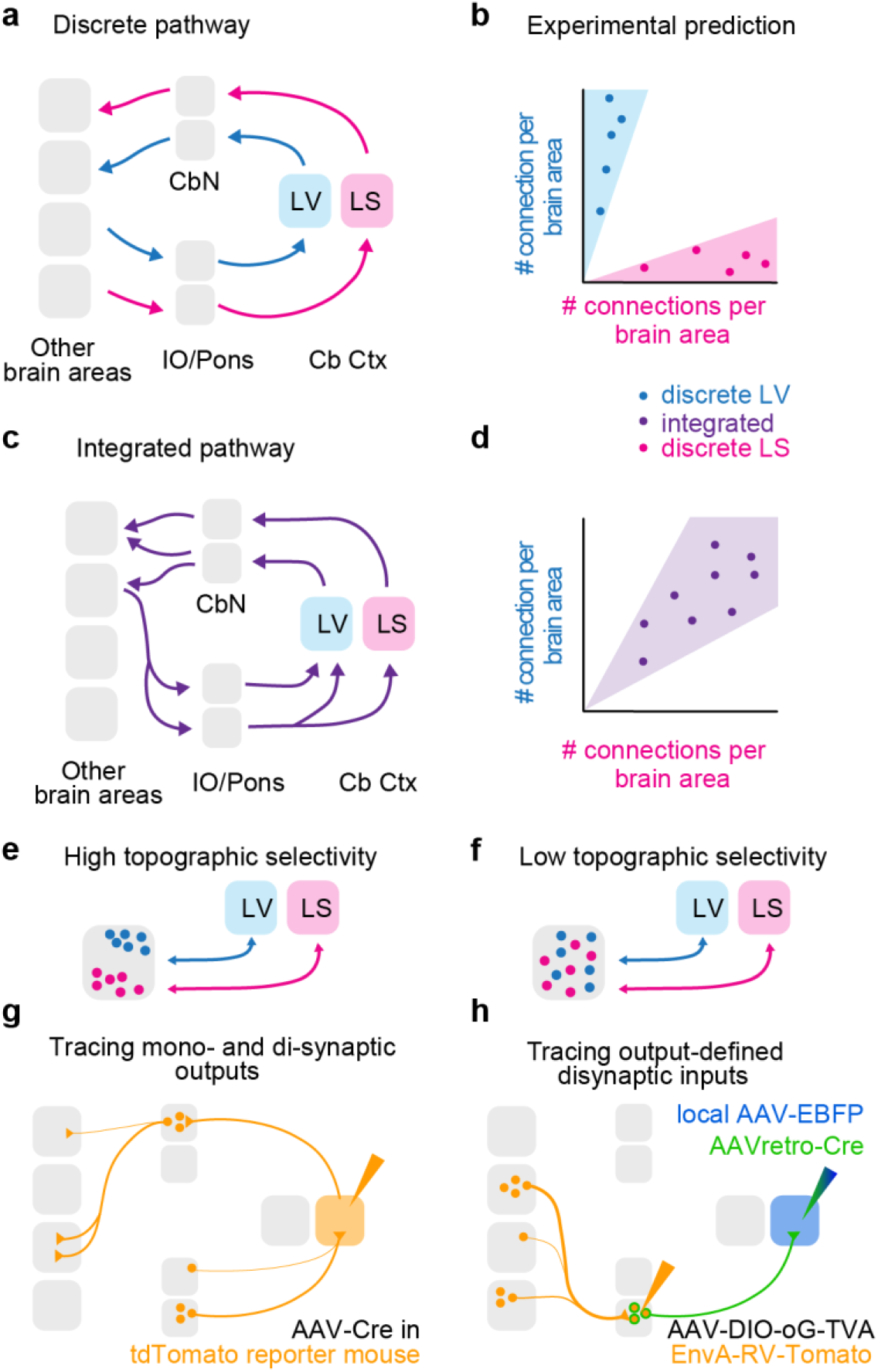
Summary of experimental approach to dissect discrete and integrated brain circuits. (**a**) Schematic representation of the discrete pathways hypothesis: LV and LS receive inputs from and send outputs to discrete sets of brain areas (**b**) The integrated pathways hypothesis: LV and LS receive inputs from and send output to integrated/shared sets of areas (**c**,**d**) Experimental prediction: discrete and integrated pathways can be revealed by comparing the connection strength between projections to LS and LV. Discrete pathways will have 10-fold stronger connections with one target; integrated pathways will have connection strength within the same order of magnitude. (**e, f**) Experimental prediction: projections to or from an area can come from spatially segregated neuronal ensembles (**e**) or from spatially intermixed neuronal ensembles (**f**). (**g**) Experimental approach to measure monosynaptic inputs and mono- and disynaptic outputs. (**h**) Experimental approach to measure disynaptic inputs via a specific monosynaptic input area.

In this framework, the spatial connection topography is a crucial determinant of what information is exchanged. Even within a single area, ensembles of neurons with different functional properties may connect to distinct targets (Kim et al., 2018). Connection patterns may be highly spatially selective if they involve spatially segregated ensembles of neurons (**Figure 1e**), or poorly spatially selective if connected ensembles are spatially overlapping (**Figure 1f**). Thus, mapping the connection topography of cerebellar lobules is necessary to identify finer scale differences in the specialization of cerebellar circuits.

Using a combination of mono- and trans-synaptic viral tracing methods, we mapped the organization and spatial topography of the long-range connections to and from LV and LS, providing a comparative map of the disynaptic circuits of these anatomically distinct but functionally related cerebellar regions. Our findings reveal a simple organizational logic: monosynaptic pathways are segregated, whereas disynaptic pathways share common brain regions but are often topographically segregated at the local level. Thus, polysynaptic cerebellar connectivity appears to be organized at the scale of local circuits rather than whole brain regions, providing a framework for understanding multiregional computations involving the cerebellum.

## Results

### A framework to map and analyze brain-wide connectivity

To map the long-range input and output pathways of LV and LS, we adopted two complementary viral tracing strategies. To map monosynaptic connections, we injected adeno-associated virus (AAV2/1) expressing Cre recombinase into either LV or LS of Cre-dependent tdTomato reporter mice (Ai9; **Figure 1g**) (Madisen et al., 2010). These injections label (1) the cell bodies of neurons projecting to this cerebellar region, via their labeled axons, (2) local neurons and their axonal projections in monosynaptic target regions, and (3) the postsynaptic partners of local neurons and their axonal projections, which were labeled by anterograde propagation of the AAV2/1-Cre virus (Zingg et al., 2017; Zingg et al., 2020). To map disynaptic inputs routed via the inferior olive, we used the TRIO approach (**Figure 1h**) (Schwarz et al., 2015). We first injected LV or LS with AAVretro or CAV viruses that infected CF axons and drove Cre-recombinase expression in olivary neurons, while concurrently injecting the inferior olive with two Cre-dependent viruses encoding the molecular toolkit for rabies tracing (TVA receptor and rabies glycoprotein G) and the enhanced green fluorescent protein (EGFP). Three weeks later, we performed a second injection of EnvA-pseudotyped ΔG rabies virus into the inferior olive, enabling selective retrograde transsynaptic labelling(Rossi et al., 2020), via expression of tdTomato, of the inputs to olivary neurons that give rise to climbing fibers targeting LV or LS. With both approaches, fixed brains were imaged using whole brain two-photon tomography to detect and trace labeled neurons and their projection patterns across the brain (**Supplementary Figure 1**).

To register, standardize, and quantify traced neurons across experiments, we developed *Braintracer*, an analysis pipeline that extends the existing *BrainGlobe* software package (Claudi et al., 2020) and its component tools (Niedworok et al., 2016; Tyson et al., 2021). Using our pipeline, we were able to: register individual brains to the Allen Common Reference atlas (Oh et al., 2014), denoise and perform background correction, normalize with a variety of measures, detect and count cells in all atlas regions, and visualize the resulting data (**Supplementary Figure 1**). For each brain, this pipeline produced an isometrically registered brain in which we could assess high-resolution axonal or somatic fluorescence with region boundaries delineated at 10 x 10 x 10 μm voxel resolution. Braintracer also augments the segmentation available in the Allen Common Reference atlas with new manual annotations of the subnuclei of the inferior olive (Luo et al., 2023) (**Supplementary Figure 2**), and a new supergroup of existing nuclei composing the mesodiencephalic junction (Wang et al., 2022) (see **Methods**).

### Segregated inputs and outputs of the cerebellar cortex

We began by assessing the degree to which CF and MF source origin and CbN target regions were similar or different between LV and LS, using experiments where we injected AAV1-Cre in TdTomato reporter mice (N = 4 for LV and LS; **Figure 2a**). We confirmed that our targeted injections labeled non-overlapping cortical territories in LV and LS by quantifying spatial patterns of labeling within the region of interest (**Figure 2b-d**). First, we computed median fluorescence signal density maps for each brain area, visualizing the density of labeling across voxels. Inspection of signal density in the cerebellar cortex revealed clearly segregated spatial patterns of labeling in the two experimental groups (**Figure 2c**). Second, to quantify this observation, we calculated the proportion of voxels in this 3D density map that exhibited 10-fold more labeling for LV than LS or vice versa (**Figure 2d**). By calculating the proportion of all voxels that exceeded this threshold, we computed a topographic selectivity index (TSI), with values closer to 1 indicating greater differential patterning between LV and LS (see **Methods**). In the cerebellar cortex, this TSI was 0.94. Targeted injections in LV and LS labeled spatially segregated inferior olive neurons (CF inputs, TSI = 0.94, **Figure 2e-h**), and more overlapping basilar pons neurons (MF inputs, TSI = 0.69) (**Figure 2i-l**). The injections also labeled segregated Purkinje cell axonal projections (and postsynaptic neurons) in the cerebellar nuclei, the monosynaptic output targets of the cerebellar cortex (TSI = 0.85, **Figure 2m-p**), consistent with a modular organization (Apps et al., 2018).

**Figure 2:**
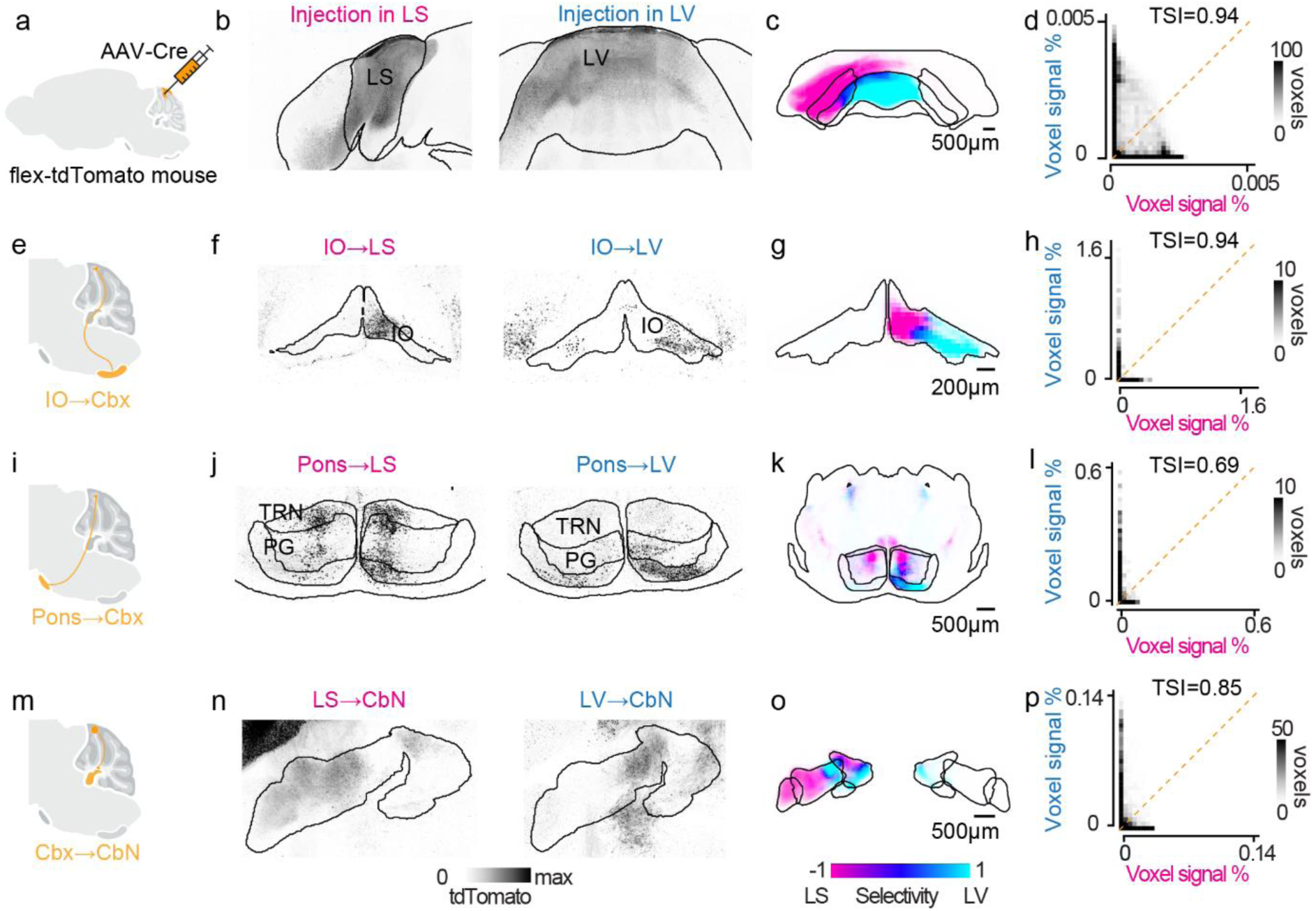
Tracing the monosynaptic inputs and outputs of LV and LS. (**a**) To label both monosynaptic inputs (CF, MF) and outputs (PCs axons) an AAV expressing Cre is injected into the target cerebellar lobule in a Cre-dependent tdTomato reporter mouse line. (**b**) Coronal brain sections through the cerebellar cortex showing tdTomato expression (black) from example injections successfully targeting LS (left) and LV (right). (**c**) Horizontal projection of the median labelling density in the cerebellar cortex for LS (n=4, magenta) and LV (n=4, cyan) injections. Data from each experiment were aligned to the Allen Common Reference atlas before averaging. (**d**) Normalized labelling density of voxels in the cerebellar cortex from experiments targeting LV (cyan, y axis) plotted against matching voxels from injections in LS (magenta, x axis). (**e**) Retrograde labelling of inferior olive (IO) neurons via AAV infection of climbing fiber axons (yellow). (**f**) Coronal section showing example IO neurons retrogradely labelled from LS and LV injections. (**g**) As in **c**, but coronal projection of labelling density in the IO. (**h**) Same as **d**, for voxels in the IO. (**i**, **j**, **k**, **l**) Same as **e**-**h**, for retrograde labelling of mossy fiber inputs from the pontine nuclei. (**m**) Purkinje output axons and anterograde transsynaptic transport of the AAV-Cre labels monosynaptically connected regions in the cerebellar nuclei. (**n**, **o**, **p**) Same as **f**, **g**, **h**, but for anterograde labelling of neurons and axons in the CbN.

We used this data to systematically identify the brain areas that either send monosynaptic efferent projections to, or receive monosynaptic afferent projections from LV or LS, and to determine differences in the distribution and the selectivity of these connections for one region or the other (**Figure 3a-d**). To identify areas with the greater differences in connection strength, we computed per area a connection selectivity index (CSI), which measures the relative difference in connection strength between each area and LV or LS (**Figure 3b**). As a reference, this analysis yielded a strong effect of experimental group (p<0.01), area (p<0.001) and their interaction (p<0.001) for the labelling at the targeted cerebellar lobules, together with high connection and topographic selectivity (CSI = 0.92 ± 0.02, TSI = 0.66 ± 0.03, mean ± s.e.m.).

**Figure 3:**
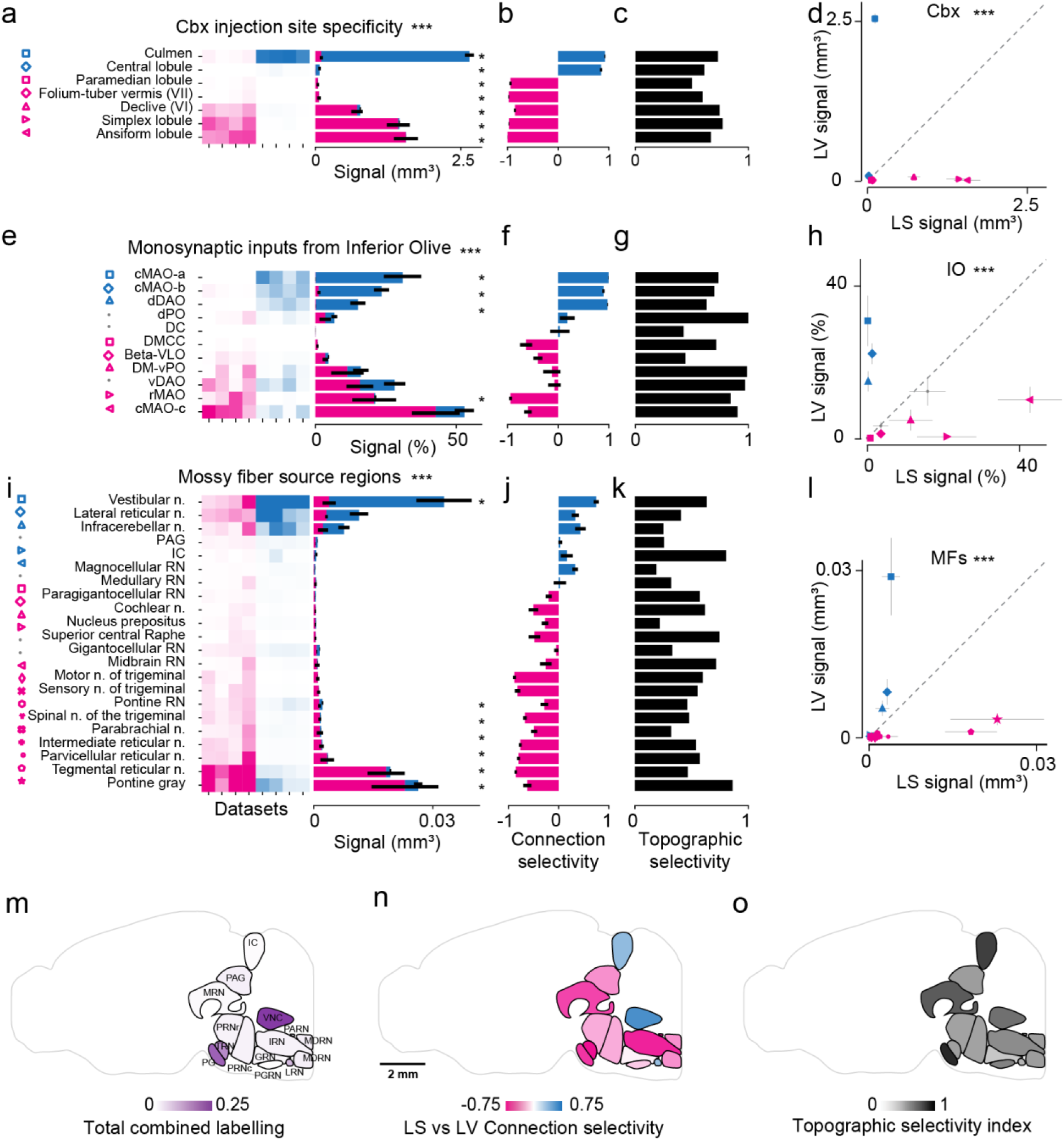
Segregated monosynaptic inputs of LV and LS. (**a**) Injection target specificity for LS and LV experimental groups. Left: Labeling intensity in relevant cerebellar cortex lobules (rows) across all experiments (columns), sorted by experimental group (LS: magenta, or LV: cyan). A two-way ANOVA reports a significant effect of area and group-by-area interaction (***, p<0.001). A follow up U-test identified areas with significantly different labelling (*, p<0.05). Right: stacked histogram of average labelling intensity across experimental groups (mean ± s.e.m. is shown for each group). (**b**) Average connection selectivity index of labelling signal for each area (mean ± s.e.m. for all pairwise comparisons between datasets across experimental groups). Negative values indicate stronger labelling in LS datasets (magenta); positive values indicate stronger labelling in LV datasets (cyan). (**c**) Topographic selectivity index for the labelling density in each area. Higher values correspond to larger differences in the spatial distribution of labelling. (**d**) Average (mean ± s.e.m.) labelling intensity in LS vs LV experimental groups for cerebellar lobules shown in **a**-**c**; areas with labeling differences greater than one standard deviation over the mean have colored markers, matching those in a. (**e, f**, **g**, **h**) Monosynaptic climbing fiber inputs from the inferior olive (IO). Same as **a-d**, for anatomical subdivisions of the IO. All subregions shown. (**i, j, k, l**) Monosynaptic mossy fiber inputs. Same as **a-d**, for nuclei providing mossy fibers, selected according to the Allen Brain Connectivity Atlas. (**m**) Areas providing monosynaptic MF inputs to the cerebellar cortex, color-coded according to normalised input strength across LS and LV datasets. A sagittal projection from the Allen CCF Atlas is shown. (**n**) Connection selectivity index for LS and LV for the monosynaptic input areas shown in **m**. (**o**) Topographic selectivity index for the input neurons within the monosynaptic input areas shown in **m.**

LV and LS received highly selective inputs from distinct sets of climbing and mossy fibers: those providing inputs to LS provided little or no input to LV and vice-versa (p <0.001 for areas; p<0.001 for group-area interaction, two-way ANOVA; CSI = 0.53 ± 0.11 for CF, 0.46 ± 0.06 for MF, mean ± s.e.m., **Figure 3e-l**). LV inputs originated from subnuclei a and b of the caudal part of the medial accessory olive (cMAO-a and cMAO-b) and the dorsal fold of the dorsal accessory olive (dDAO, p<0.05, U-test), while LS received CF input primarily from subnucleus c of the caudal part of the medial accessory olive (cMAO), rostral part (rMAO, p<0.05, U-test) and the ventral fold of the dorsal accessory olive (vDAO) (**Figure 3e**). Moreover, LV received more abundant MF inputs from the vestibular (p<0.05, U-test), lateral reticular and infracerebellar nuclei, while LS inputs originated more selectively from the pontine gray (p<0.05, U-test) and the tegmental reticular nucleus (p<0.05, U-test) (**Figure 3i**). In regions that projected to both LV and LS, projections often arose from non-overlapping parts of these structures (CF TSI = 0.76 ± 0.06, MF TSI = 0.50 ± 0.04, mean ± s.e.m., **Figure 3g,k)**, possibly reflecting nuclei, layering, or domains not annotated in the standard atlas (see **Methods** for area selection algorithm). Specifically, in line with previous anatomical studies (Apps & Hawkes, 2009; Luo et al., 2023), CF projections to LV in the vermis arose predominantly from the caudal part of the medial accessory olivary nucleus, whereas projections to LS in the intermediate hemispheres arose from both the medial and dorsal accessory olivary nuclei (**Figure 2g** and **Figure 3e**). Within the pons, which receives topographically organized cortical inputs (Henschke & Pakan, 2020; Leergaard & Bjaalie, 2007), the tegmental reticular nucleus (TRN) and the pontine gray displayed significant spatial segregation of inputs (combined TSI = 0.85, **Figure 3i** and **Supplementary Figure 3s,t**). Regions such as the periaqueductal gray (PAG) had a low topographic selectivity index, indicating an overlapping spatial distribution of labeling (TSI = 0.26, **Figure 3i, Supplementary Figure 3c,d**).

Similarly to afferent inputs, Purkinje cells in LV and LS exhibited non-overlapping monosynaptic efferent projections to the cerebellar nuclei (p <0.001 for areas, p<0.01 for group-area interaction, two-way ANOVA, CSI = 0.65 ± 0.11, mean ± s.e.m., **Figure 2n-p** and **Figure 4**). Purkinje cells in LV did not project to the dentate nucleus (p<0.05, U-test), while LS Purkinje cells sent very sparse projections to the vestibulocerebellar nucleus (**Figure 4a-b, d-f**). In the nuclei that received input from both cortical lobules (fastigial and interposed), the spatial pattern of projections was highly non-overlapping (TSI = 0.68 ± 0.08, **Figure 4c,g**). The vestibulocerebellar nucleus was the only nucleus to show denser connectivity with LV than LS (i.e. selective for LV).

**Figure 4:**
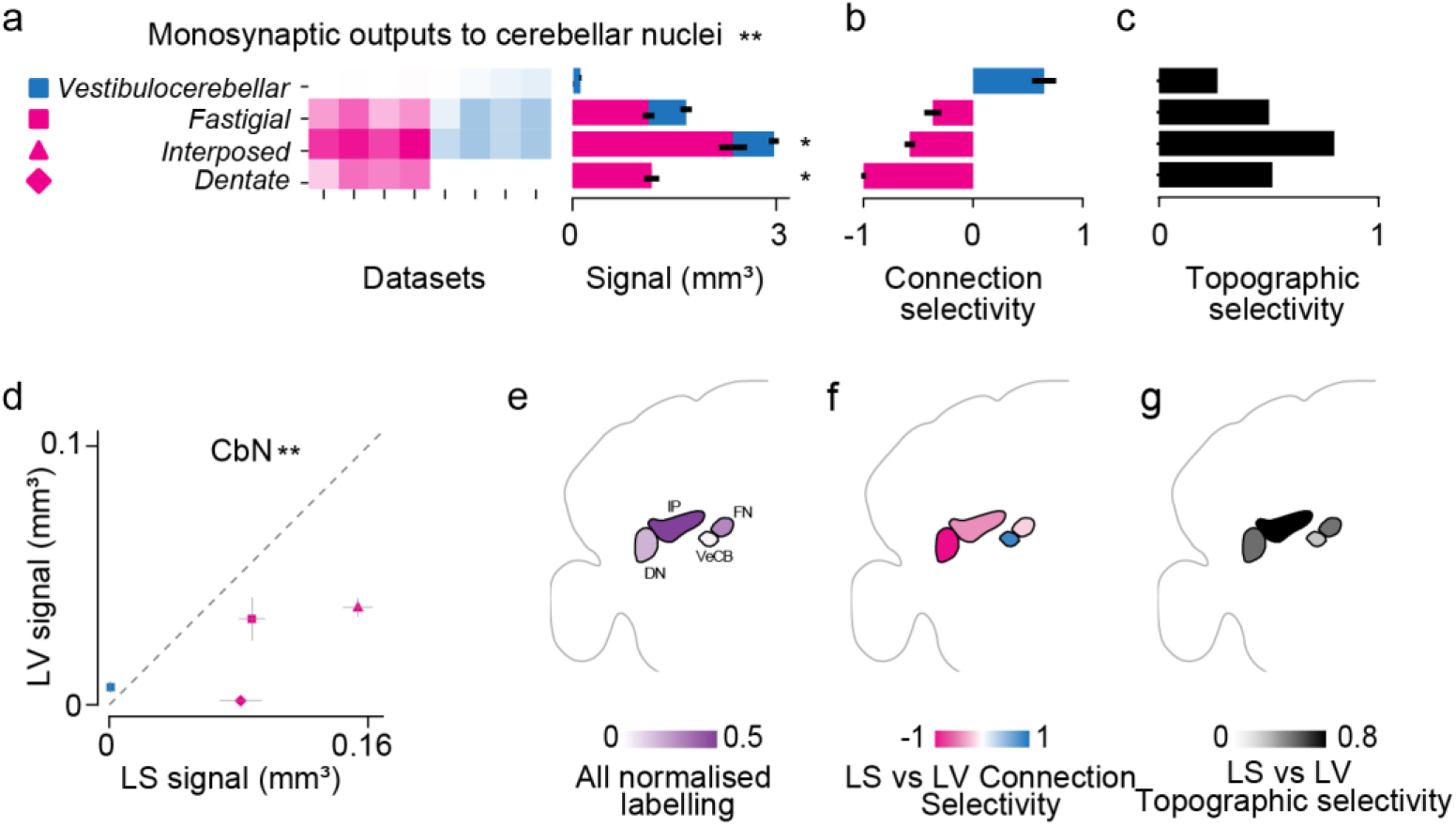
The monosynaptic outputs to the cerebellar nuclei (CbN). (**a**) Left: Labeling intensity in cerebellar nuclei (rows) across all experiments (columns), sorted by experimental group (LS: magenta, or LV: cyan). A two-way ANOVA reports a significant effect of area and group-by-area interaction (**, p<0.01). A follow up U-test identified areas with significantly different labelling (*, p<0.05) Right: stacked histogram of average labelling intensity across experimental groups (mean ± s.e.m. is shown for each group). (**b**) Average connection selectivity index of labeling signal for each nucleus (mean ± s.e.m. for all pairwise comparisons between datasets across experimental groups). Negative values indicate stronger labelling in LS datasets (magenta); positive values indicate stronger labelling in LV datasets (cyan). (**c**) Topographic selectivity index for the labelling density in each nucleus. Higher values correspond to larger differences in the spatial distribution of labelling. (**d**) Average (mean ± std) labelling intensity in LS vs LV experimental groups for output nuclei shown in **a**-**c**, where areas with labeling difference greater than one standard deviation over the mean have colored markers, matching those from a. (**e**) Areas receiving monosynaptic outputs from the cerebellar cortex, color-coded according to normalised output strength across LS and LV datasets. A coronal projection from the Allen CCF Atlas is shown. (**f**) Connection selectivity index for LS and LV to the monosynaptic output areas shown in **e**. (**g**) Topographic selectivity index of the projections within the monosynaptic output areas shown in **e.**

Taken together, these results demonstrate that the monosynaptic CF and MF afferents, as well as the monosynaptic efferents of Purkinje cells in the CbN, are highly segregated between LV and LS.

### Integrated disynaptic inputs via the Inferior Olive

Given the strong selectivity of olivary CF projections (**Figure 3e)**, we asked if the brain-wide patterns of inputs to these ensembles maintained a similar logic of segregation (**Figure 5)**. To identify the inputs to olivary neurons projecting to LV and LS, we utilized the TRIO method for input-output tracing (**Figure 5a**; (Schwarz et al., 2015); see **Methods**). In a subset of experiments, we confirmed accurate lobule targeting by co-injecting a supplementary AAV expressing EBFP in local neurons (**Figure 5b**). The topographic quantification of EBFP labeling at the injection sites confirmed that we obtained good separation of viral infection spread between LV and LS within the cerebellar cortex (TSI = 0.99, **Figure 5c,d**), better than that achieved in monosynaptic tracing experiments (**Figure 2d)**. Further confirming consistent targeting across experiments, the ensembles of first-order presynaptic CF olivary neurons projecting to LV and LS were topographically segregated and qualitatively resembled those resulting from the monosynaptic tracing experiment (regional TSI = 0.85, **Figure 5e,f**, **Figure 2f,g**). Some differences in labelling with the former strategy were expected due to the smaller injections, and since we counted both neurons expressing EGFP and tdTomato, i.e. starter cells, or tdTomato alone, i.e. potential local disynaptic inputs or starter cells with weak EGFP expression (**Figure 3e, Supplementary Figure 4)**.

**Figure 5:**
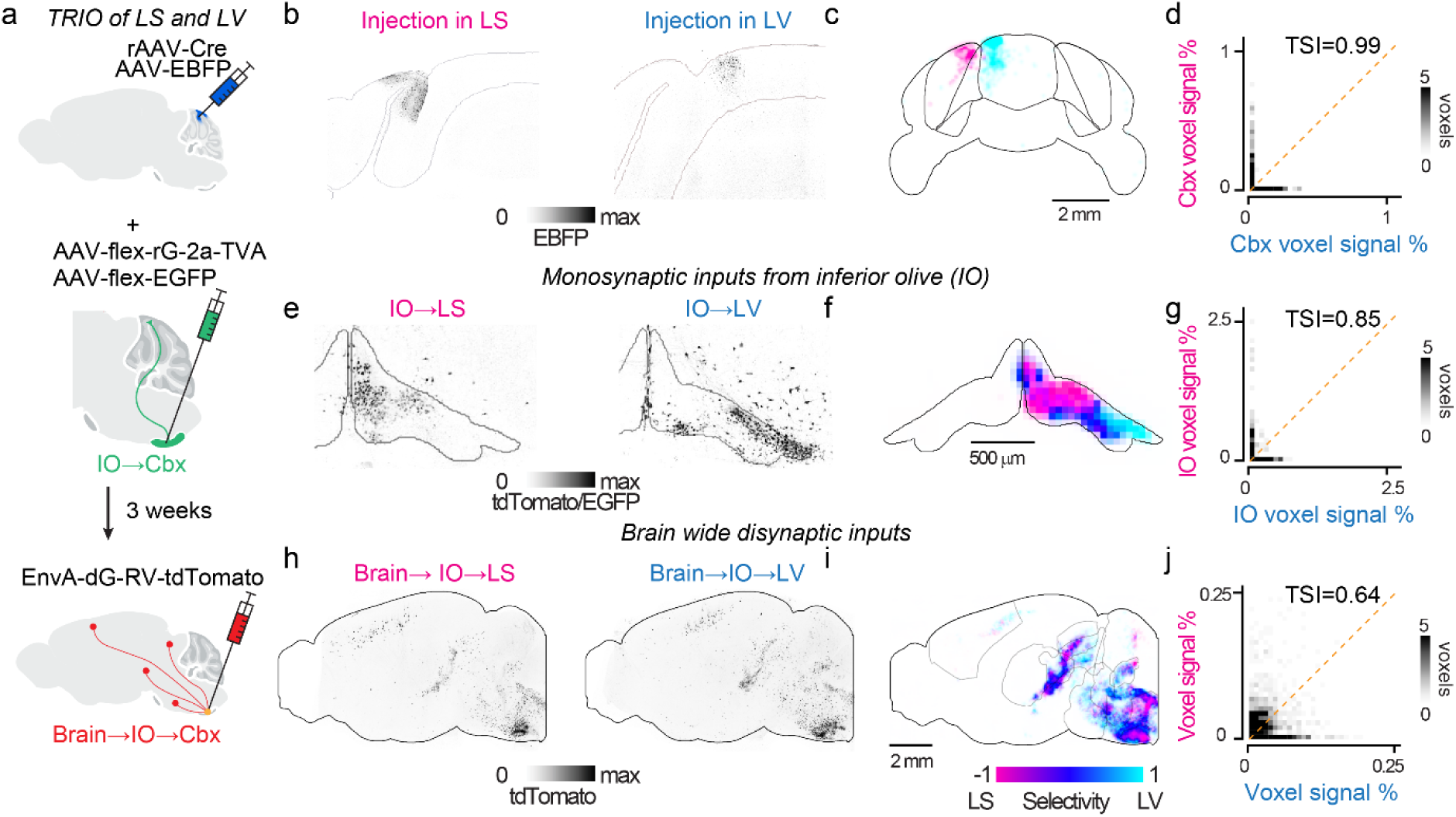
Tracing the disynaptic inputs to LS and LV via the IO. (**a**) Schematic of TRIO from the cerebellar cortex. Top: a mix of retrograde AAV expressing Cre (rAAV-Cre) and a local AAV expressing a blue fluorophore (AAV-EBFP) is injected in the target lobule of the cerebellar cortex. Middle: next, a mix of Cre-dependent AAVs expressing the rabies tracing toolkit (rG and TVA) and a green fluorophore (EGFP) are injected in the IO. The retrograde transport of the rAAV labels along climbing fiber axons triggers expression of EGFP (green) and rabies genes in monosynaptically connected inputs from the inferior olive (IO). Bottom: Three weeks later, an EnvA-dG-TdTomato rabies virus is injected in the IO, where it can only infect the neurons projecting to the cortical injection site (yellow), and then propagate to their brain-wide synaptic inputs. (**b**) Coronal sections through the cerebellar cortex demonstrating successful targeting of LS (left) and LV (right). (**c**) Coronal projection of the median labelling density in the cerebellar cortex for tracing experiments targeting LS (n=3, magenta) and LV (n=2, cyan). Data from each experiment were aligned to the Allen Common Reference atlas before averaging. (**d**) The normalised labelling density (coronal projection) from voxels in the cerebellar cortex from experiment targeting LV (cyan, y axis) plotted against matching voxels from injections in LS (magenta, x axis). (**e**) Same as **b**, for monosynaptic input CFs. (**f**) Same as **c**, coronal projection for labelling density in the IO for tracing experiments targeting LS (n=8, magenta) and LV (n=9, cyan). (**g**) Same as **d**, for voxels in the IO. (**h, i, j**) Same as **e**-**g**, for brain-wide disynaptic inputs (sagittal projection).

We labelled and traced thousands of disynaptic neurons across brain areas in the pons, midbrain and cerebral cortex (**Figure 5e**, **Figure 6**). As expected from rabies transsynaptic spread (Tran-Van-Minh et al., 2023), the number of traced neurons scaled proportionally with the number of postsynaptic olivary neurons labelled (**Figure 6e**). Approximating this relationship with a linear model (R = 0.53, p<0.05), we estimated a yield of 7.6 brain-wide inputs traced per olivary neuron (n = 16, combined across experimental groups).

**Figure 6:**
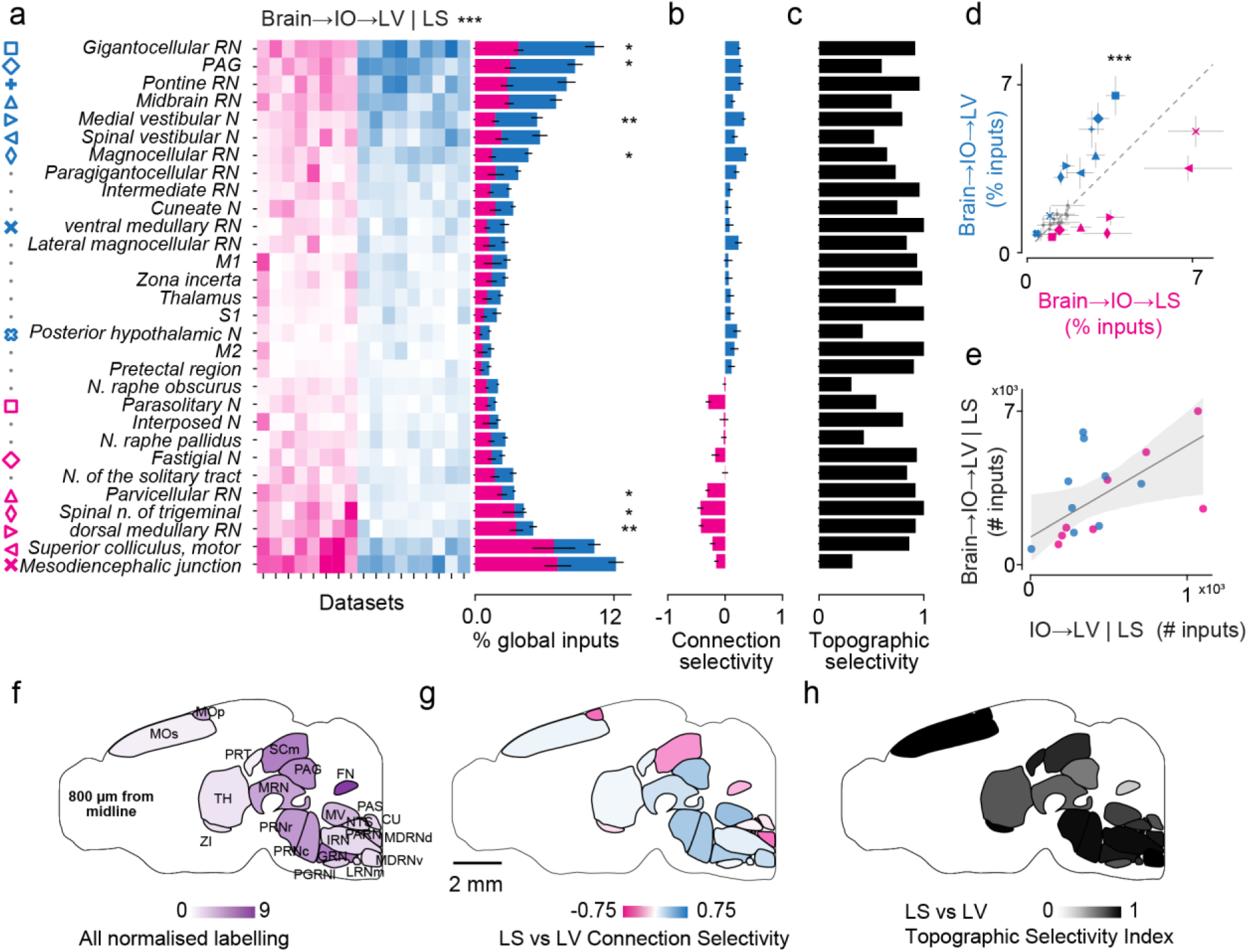
Shared disynaptic inputs to LV and LS via the inferior olive (IO). (**a**) Distribution of LV- and LS-projecting disynaptic inputs via the IO, from selected brain areas across datasets (column), sorted by experimental group (LS: magenta, or LV: cyan). Areas chosen by non-overlapping atlas hierarchy search algorithm based on significant labeling. A two-way ANOVA reports a significant effect of area and group-by-area interaction (***, p<0.001). A follow up U-test identified areas with significantly different labelling (*, p<0.05, ** p<0.01) Right: stacked histogram of average normalized disynaptic input neurons across experimental groups (mean ± s.e.m. is shown for each group). (**b**) Average connection selectivity index of the input strength from each area (mean ± s.e.m. for all pairwise comparisons between datasets across experimental groups). Negative values indicate stronger labelling in LS datasets (magenta); positive values indicate stronger labelling in LV datasets (cyan). (**c**) Topographic selectivity index for the labelling density in each upstream area. Higher values correspond to larger differences in the spatial distribution of labelling. (**d**) Average (mean ± s.e.m.) labelling intensity in LS vs LV experimental groups for brain areas in **a**-**c**, where areas with labeling differences greater than one standard deviation over the mean have colored markers, matching those in a. (**e**) Number of traced brain-wide disynaptic inputs plotted against the number of olivary neurons labeled in each dataset (LS magenta; LV cyan). Linear fit (gray line, R= 0.53, p <0.05) with confidence intervals (shaded area). (**f**) Areas providing disynaptic inputs to the cerebellar cortex via the inferior olive, color-coded according to fraction of projecting neurons traced across LS and LV datasets. A sagittal projection from the Allen CCF Atlas is shown. (**g**) Connection selectivity index for LS and LV for the disynaptic input areas shown in **g**. (**h**) Topographic selectivity index for the traced neurons within the disynaptic input areas shown in **g**.

While the overall brain-wide connection patterns differed between LV and LS (**Figure 6a,d**, p<0.001 for areas; p<0.001 for group-area interaction, two-way ANOVA), there was notably more overlap and lower connection selectivity in the disynaptic input brain regions than in their postsynaptic ensembles in the IO (p<0.01, Kolmogorov-Smirnov [K-S] test, **Figure 3f**, **Figure 6b**). The greater differences in connection patterns were driven by the gigantocellular reticular nucleus (p<0.05, U-test), the PAG (p<0.05, U-test), the medial vestibular nucleus (p<0.01, U-test), the magnocellular reticular nucleus (p<0.01, U-test), the parvicellular reticular nucleus (p<0.05, U-test), spinal nucleus of the trigeminal (p<0.05, U-test), and the dorsal medullary reticular nucleus (p<0.01, U-test). The motor superior colliculus, the pontine reticular nucleus and the mesodiencephalic junction also provided dominant input, but did not show significant differences in connection strength between LS and LV.

While connection selectivity was low (CSI = 0.18 ± 0.02, mean ± s.e.m. across regions), the analysis of topographic distribution of source neurons *within* a given region revealed that input neurons often resided in locally non-overlapping territories (**Figure 6c**, average TSI = 0.77 ± 0.04, mean ± s.e.m. across regions). We analyzed the topographic organization of neuronal inputs from several of these inferior olive input brain regions, including the somatomotor cortex and mesodiencephalic junction (MDJ), in greater detail (**Figure 6f-h, Supplementary Figure 6**). MDJ inputs to LV and LS were largely overlapping (TSI = 0.32), while those from the somatomotor cortex, thalamus, pontine reticular nuclei, cerebellar nuclei, and the motor layer of the superior colliculus demonstrated marked distinction in the spatial origin of their inputs to these cerebellar lobules (**Figure 6f-h**, TSI ranging from 0.73-0.93).

Taken together, our TRIO tracing of inputs to olivary neurons that project CFs to LV and LS demonstrates a fundamental organizing principle. Olivary neurons projecting to the cerebellar cortex are highly segregated but receive input from a common set of forebrain, midbrain, and hindbrain regions (**Figure 6f, g)**. However, the neurons within each of these source regions that target LV and LS projecting olivary regions are likely to be independent (**Figure 6h)**. Thus, the information streams of instructive signals sent to the cerebellar cortex, while shared across brain regions, are topographically specified at microcircuit level.

### Absence of dopaminergic input from the ventral tegmental area to the inferior olive

Our TRIO experiments did not reveal inputs to the IO from the ventral tegmental area (VTA), a candidate source region for driving the reward signals that have been recorded in climbing fibers in LS and, to a lesser extent, in LV (Heffley & Hull, 2019; Kostadinov et al., 2019; Kostadinov et al., 2025). A projection from VTA to IO has been previously reported using non-specific bulk labelling of VTA neurons projecting to the dorsal cap of the inferior olive (a region sending very few projections to LV and LS; **Figure 3e**) (Fallon & Bañales, 1984). We therefore performed further experiments to confirm that dopaminergic VTA projections were not missed due to potential neural tropism of the TRIO viruses.

We labelled the brain-wide axonal projections of dopaminergic neurons by injecting an AAV2/1-FLEX-tdTomato virus into the VTA of DAT-iCre mice (N = 3). This produced strong labeling throughout the midbrain dopamine system but no detectable axonal fluorescence in the IO (**Supplementary Figure 7**). To increase the sensitivity of this assay for detecting fine axonal terminations, we imaged samples at the higher spatial resolution of 1x1x5 μm per voxel. These analyses did not identify any difference in IO labelling between injected DAT-Cre mice and wild-type controls (**Supplementary Figure 7**). This provides anatomical support for the idea that reward signals in climbing fibers are not directly inherited from dopaminergic neurons in the VTA.

### The disynaptic outputs of the cerebellar cortex

Finally, we sought to determine the downstream targets of the cerebellar nuclei neurons that receive the output of the LS and LV pathways (**Figure 7**). To map these disynaptic outputs, we took advantage of the anterograde transsynaptic properties of the AAV2/1-Cre virus that we employed to map the monosynaptic circuits (Zingg et al., 2017; Zingg et al., 2020). We reasoned that injections in the cerebellar cortex would result not only in the labeling of the output axonal tracts of Purkinje neurons, but also in the trans-synaptic transduction of postsynaptic cerebellar nuclei neurons and their axonal projection to disynaptic output areas.

**Figure 7:**
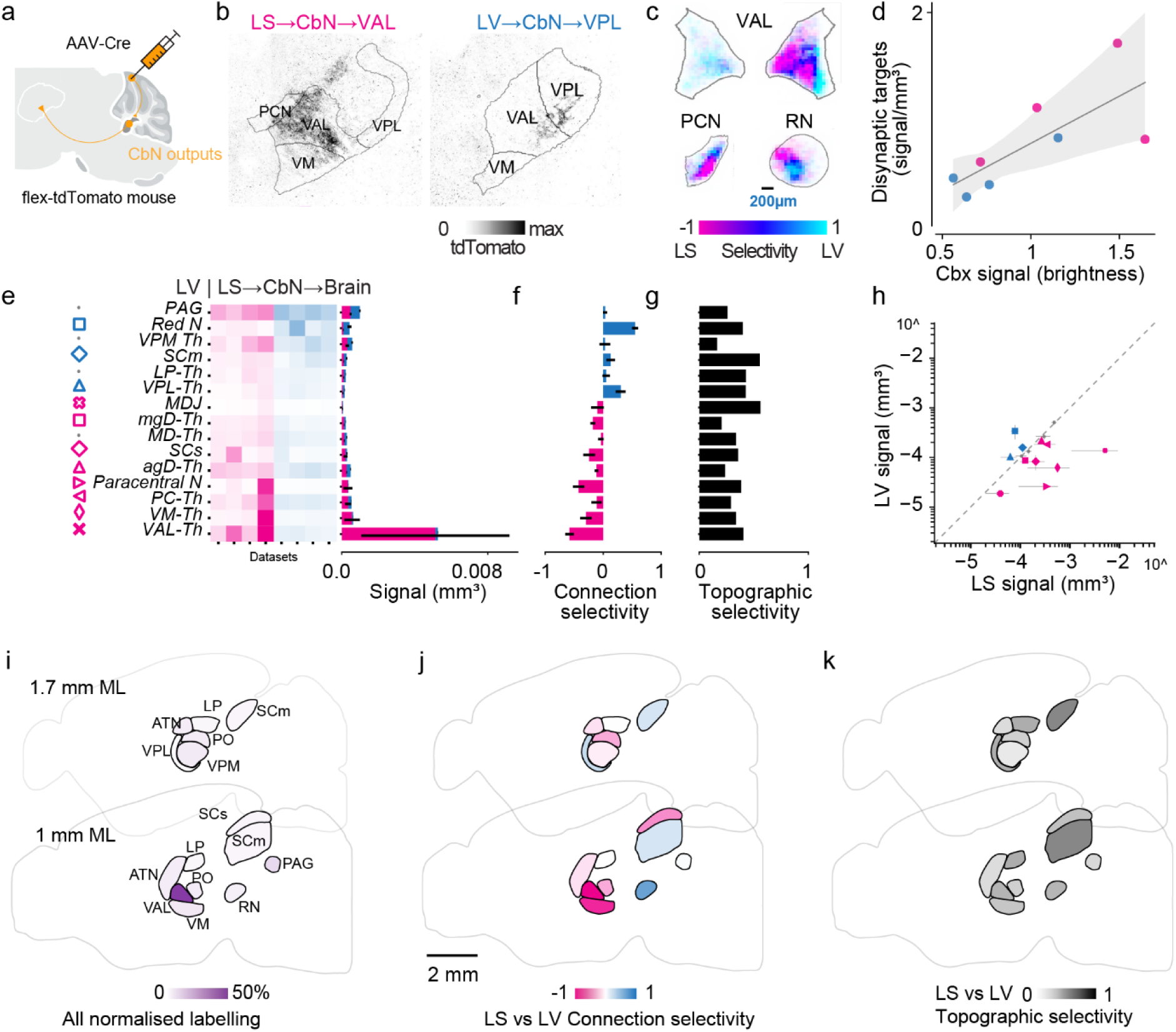
Integrated disynaptic outputs of LS and LV. (**a**) Schematic of anterograde transsynaptic tracing. Following injection of an AAV expressing Cre (yellow syringe) into the cerebellar lobule of interest in a Cre-dependent tdTomato reporter mouse line (cfr. Fig 2), the anterograde monosynaptic transport of the AAV-Cre infects postsynaptic output neurons in the cerebellar nuclei (yellow dot); the axonal projections of these neurons (yellow) are labelled throughout the brain. (**b**) Example axonal projection (maximum) labelling in the ventral anterior lateral nucleus of the thalamus (VAL) from an injection in LS (left) and in VAL and ventral posterolateral nucleus (VPL) from an injection in LV (right). (**c**) Coronal projection of the average labelling density in the VAL, paracentral nucleus (PCN), and red nucleus (RN) for tracing experiments targeting LS (n=4, magenta) and LV (n=4, cyan). Data from each experiment were aligned to the Allen Common Reference atlas before averaging. (**d**) Total signal volume in downstream disynaptic target regions plotted against normalized cerebellar cortical signal brightness in the original image stack for each dataset (LS magenta; LV cyan). Linear fit (gray line, R= 0.80, p <0.05) with confidence intervals (shaded area) (**e**) Left: disynaptic axonal labelling volume from an injection in LS (magenta) or LV (cyan) from selected brain areas (rows) in each dataset (column), sorted by experimental group (LS: magenta, LV:cyan). A two-way ANOVA reported no significant effect of area and group-by-area interaction. Right: stacked histogram of average axonal labelling volume across areas and experimental groups (mean ± s.e.m. is shown for each group). (**f**) Average connection selectivity index of the axonal labeling in each area (mean ± s.e.m. for all pairwise comparisons between datasets across experimental groups). Negative values indicate stronger projections from LS (magenta); positive values indicate stronger projections from LV (cyan). (**g**) Topographic selectivity index for the axonal distribution in each area. Higher values correspond to larger differences in the spatial distribution of axons. (**h**) Average (mean ± std) volume of disynaptic axonal labeling in LS vs LV experimental groups for areas shown in **e**-**g**, where areas with labeling differences greater than one standard deviation over the mean have colored markers, matching those in e. (**i**) Areas receiving disynaptic outputs from the cerebellar cortex, color-coded according to normalized output strength across LS and LV datasets. Two sagittal projections from the Allen CCF Atlas are shown, 1mm and 1.7mm medio-lateral from bregma. (**j**) Connection selectivity index for LS and LV for the disynaptic output areas shown in **i**. (**k**) Topographic selectivity index for the projections traced within the disynaptic output areas shown in **i**.

Indeed, we were able to detect fluorescent labelling of both somas in cerebellar nuclei (**Figure 2n**), their decussating projections, and axonal arborization in downstream targets such as the thalamus and red nucleus (**Figure 7a-c**). To avoid contamination of this analysis with directly labeled monosynaptic input neurons (**Figure 2**, **Figure 3**), we restricted our analysis to brain areas that are known to receive projections from the cerebellar nuclei (see **Methods** for list). As expected, the amount of these axonal projections in these areas scaled linearly with the labeling at the cerebellar cortex injection site (R = 0.80, p <0.05, **Figure 7d**).

The major output targets were the thalamus, the superior colliculus, the PAG, and the red nucleus: these outputs were largely overlapping (p>0.5 for all factors, two-way ANOVA), with low connection selectivity (CSI = 0.21 ± 0.05) and topographic selectivity (TSI = 0.35 ± 0.03). Albeit not significant, the strongest differences in projections patterns among the thalamic nuclei were observed in the ventral anterior-lateral nucleus (VAL, LS) and in the ventral posterolateral nucleus (VPL; LV) (**Figure 7e-f**). We also found strong labelling in areas such as the ventromedial nucleus (VM) from a subset of LS datasets, but this appeared to form part of a broader connection with VAL (**Figure 7b**). The preferential connectivity of this area with LS aligns with results from previous TRIO experiments involving VM and the fastigial nuclei (Zhu et al., 2023). The red nucleus demonstrated significant connection selectivity for LV. We also found that the MDJ does not serve as a cerebellar output from LV or LS (**Figure 7e**).

These results highlight a topographic mapping of cerebellar modules onto the thalamic relay that is less stark than in monosynaptic input/output regions, but nonetheless could enable differential output to the rest of the brain (**Figure 7i-k**).

### Diverse architectures for cerebellar computation

Our data suggest that the monosynaptic and disynaptic pathways of LV and LS follow different connectivity logic. Monosynaptic connections are segregated, with pronounced differences in the regional origins of climbing fiber and mossy fiber inputs (**Figure 3**), as well as distinct projection targets within the cerebellar nuclei (**Figure 4**). In contrast, disynaptic connections are more integrated: input and target neurons belong to overlapping ensembles of areas that often differ in fine-scale spatial topography within those areas (**Figure 6**, **Figure 7**).

To quantitatively compare the degree of segregation and local topography of these pathways, we plotted the LV vs LS connection selectivity index against the within-regional spatial selectivity index for brain areas significantly contributing to monosynaptic and disynaptic circuits of LV and LS (**Figure 8**). Areas providing disynaptic inputs to cerebellar lobules formed a cluster with significantly lower connection selectivity (p<0.001, K-S test, **Figure 8a**), but significantly higher spatial selectivity than areas providing monosynaptic inputs (p<0.001, K-S test, **Figure 8a**). This supports a connection logic whereby pathways originating from different local circuits in an overlapping set of brain areas converge onto distinct monosynaptic cerebellar circuits, especially in the climbing fiber input pathway (**Figure 8b**).

**Figure 8:**
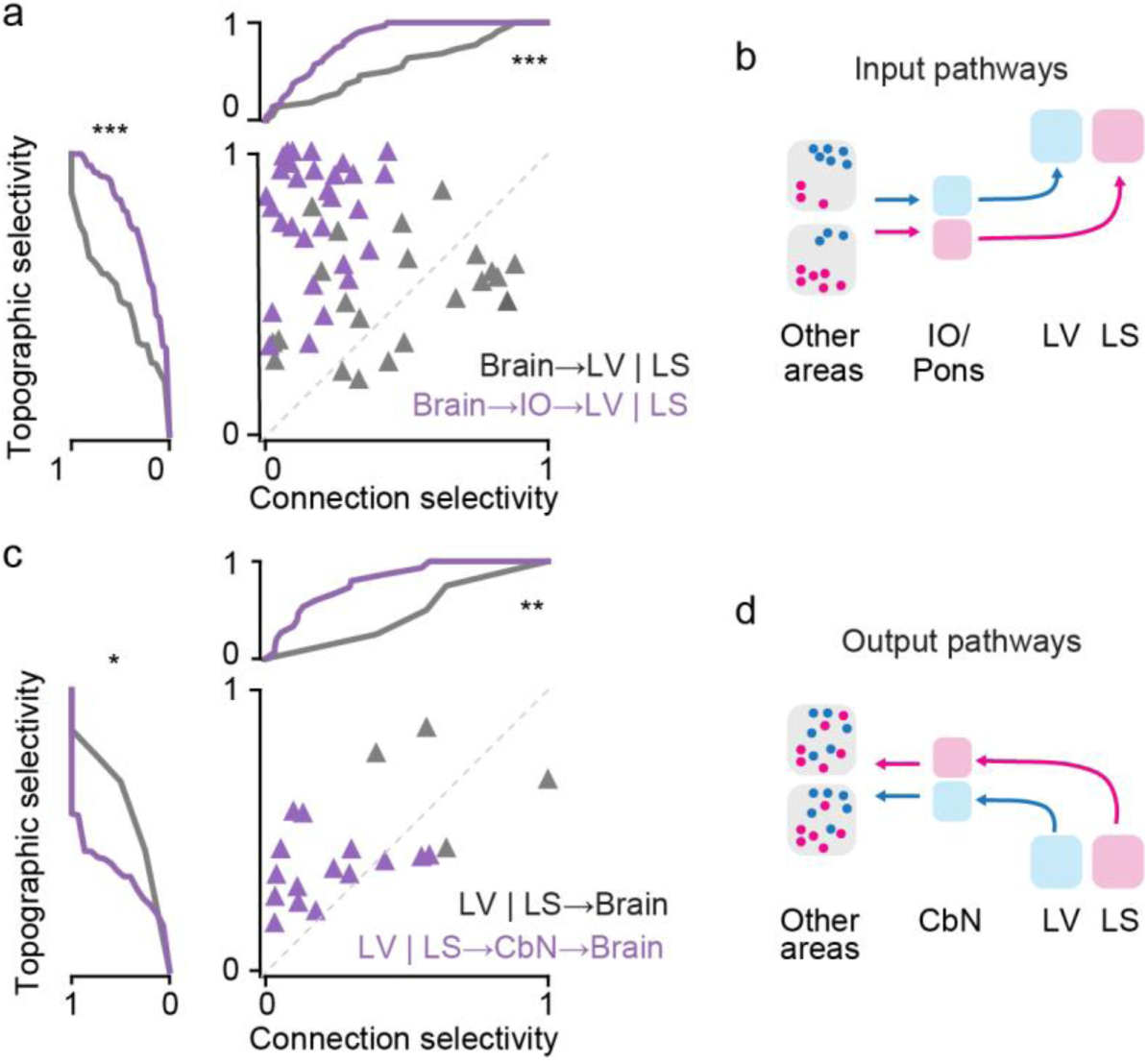
Segregated and integrated circuits of the cerebellar cortex. (**a**) The topographic selectivity index of mono- (gray) and disynaptic (purple) input areas plotted against their connection selectivity index for LS or LV. Marginal cumulative distributions are shown (p<0.001 for selectivity index, p<0.001 for topographic selectivity, K-S test). (**b**) Schematic of proposed circuit logic of input connections to LV and LS. Disynaptic inputs from various brain areas tend to have lower connection selectivity and higher topographic selectivity than monosynaptic inputs from the IO or the pons. (**c**) Same as **a**, for mono- (orange) and disynaptic (purple) output areas. (p<0.05 for selectivity index, p<0.01 for topographic selectivity, K-S test) (**d**) Same as **b**, for the output connections from LV and LS. Disynaptic outputs to other brain areas tend to have both lower connection selectivity and lower topographic selectivity than monosynaptic outputs to the cerebellar nuclei.

We then repeated the same analysis for areas receiving outputs from the cerebellar lobules: monosynaptic targets formed a cluster with significantly higher connection selectivity (p<0.05, K-S test, **Figure 8c**), and significantly higher spatial selectivity than areas receiving disynaptic outputs (p<0.01, K-S test, **Figure 8c**). This supports a connection logic whereby the output of each cerebellar lobule is relayed from distinct monosynaptic output pathways to overlapping circuits and brain areas (**Figure 8d**).

## Discussion

The canonical architecture of the cerebellar microcircuit has led to the prevailing idea that a common neural computation is performed across cerebellar cortical modules. This conserved local structure is in stark contrast to the long-range connectivity of different parts of the cerebellum, which likely form diverse polysynaptic loops with cortical and subcortical partner regions involved in a variety of motor and cognitive behaviours. These differences in long-range connectivity may recruit the cerebellar canonical circuit in different lobules to serve lobule- or module-specific functions.

Here, we set out to determine the anatomical underpinnings of the encoding properties of lobules V (LV) and simplex (LS), cerebellar regions that, while being both involved in processing forelimb movements, exhibit complementary encoding of movement and reward (Kostadinov et al., 2025). By using a combination of AAV2/1 and rabies-mediated viral tracing methods, we mapped the local and long-range organization of the extracerebellar connections formed between these regions and the rest of the brain.

### Comparative organization of monosynaptic and disynaptic connections

At the monosynaptic level, LV and LS share very few input and projection targets (**Figure 3**, **Figure 4**): they receive CF input from distinct subregions of the inferior olive, MF input from distinct mossy fiber source regions, and send projections to distinct parts of the cerebellar nuclei, consistent with a modular organization of these pathways (Apps et al., 2018). In contrast to this segregated monosynaptic connectivity, LV and LS receive disynaptic input via the inferior olive from a shared set of brain-wide regions: those providing significant input to either lobule also project to the other lobule. (**Figure 6**). Differences emerge only at the level of patterns of connection volume, and more strongly in the topographic distribution of individual connected neurons within these regions (**Figure 6c**). This suggests that LV and LS receive disynaptic input from segregated populations of neurons in overlapping sets of brain regions, configuring private lines of information processing from the same brain areas serving parallel computations in the cerebellar cortex (Kostadinov et al., 2025). Disynaptic output regions exhibit even greater mixing: targets of the cerebellar nuclei that are innervated by LV and LS Purkinje cells lose both connection and spatial selectivity compared to the cerebellar nuclei themselves. This suggests that the computations performed by these circuits can be used by a similar set of long-range downstream partners. Thus, the cerebellum may perform computations using parcellated instructive signals for different features of an action or thought and then emit an integrative output signal to its target regions to affect their neural dynamics (Li & Mrsic-Flogel, 2020). In this framework, LV and LS could compute motor error or reward-based metrics for the control of forelimb movements based on differential input features; their output would be unified in a single forelimb control signal in downstream regions.

### Computational implications of differences in Lobule V and simplex connectivity

LV and LS are both implicated in forelimb motor control but have complementary encoding properties: LV preferentially exhibits movement-related climbing fiber signals while LS preferentially exhibits reward-related signals (Kostadinov et al., 2025). The local differences in disynaptic input source neurons within similar parent regions may be the substrate for this complementary encoding.

This sensorimotor organization may reflect the structure of inputs to the inferior olive from motor cortical regions, which reach the olive either directly or indirectly via the mesodiencephalic junction (MDJ). Although direct projections from motor cortex are relatively sparse, they are highly segregated at the local scale and may therefore convey distinct streams of functional information. In contrast, the more numerous projections from the MDJ show greater overlap. Neurons in the MDJ are known to receive smooth cortical projection gradient (Wang et al., 2022), suggesting that they can relay integrated information across modalities. Together, these disynaptic pathways could convey related functional streams to LV and LS, allowing different movement-related features—for example, shoulder control, wrist control, or combinations of these variables—to be processed separately within the cerebellum.

The origin of reward-related climbing fiber signals is less clear, as we did not detect projections from the midbrain dopamine system to the cerebellum via the inferior olive. If the reward signals recorded in LS climbing fibers are not dopaminergic in origin, their source remains to be determined. Our anatomical data argue specifically against a direct dopaminergic VTA→IO projection as the source of reward-related climbing fiber signals, and non-dopaminergic VTA projections to olivary subregions outside the LV/LS climbing-fiber territory (Fallon & Bañales, 1984; Winship et al., 2006). Since reward signals in granule cells are also not inherited directly from the dopamine system (Wagner et al., 2019), one intriguing possibility is that reward-related signals throughout the cerebellum arise from polysynaptic routes through the cortex or MDJ. Candidate pathways include direct inputs from somatomotor cortex, and potentially also from prefrontal or orbitofrontal cortex, where we traced a small number of neurons in a subset of TRIO experiments targeting LS (**Figure 5h**). Alternatively, cortical reward signals may reach the cerebellum through indirect connections via the multimodal neuronal ensembles in the MDJ (Wang et al., 2022).

In contrast to input pathways to LV and LS, cerebellar outputs from these regions show a higher degree of convergence in the thalamus (Dacre et al., 2021), suggesting they may be utilized collectively, including when they are further routed downstream to motor cortical targets. Thus, the outputs of distinct cerebellar internal model computations (Ito, 2008; Wolpert et al., 1998) may be integrated to modify dynamics and learn (Gao et al., 2018; Li & Mrsic-Flogel, 2020; Zhu et al., 2023).

### Utility of our dataset and tool

We have developed and release *Braintracer*: a complete processing pipeline for anatomical neuronal tracing. The Python package can easily be installed from the command line with PyPI (https://pypi.org/project/braintracer/). Users have access to the *Braintracer* wrapper of Cellfinder (Tyson et al., 2021), which enables easier pre-processing of datasets for analysis (atlas registration and cell detection) and provides tools for further training of the 3D ResNet convolutional neural network for cell detection. It also provides an optional pre-processing pipeline that detects signal volume rather than somas, producing output in the same format. The bulk of *Braintracer* consists of a suite of visualization options for 3D anatomical images and tracing data, such as those generated by Cellfinder. Crucially, it provides an easy way to compare results across many datasets and between two groups. The majority of plots shown in this paper are outputs of this suite and can be flexibly adapted to display other users’ data. We are providing this package for open access under a GPLv3 license (see **Methods**).

### Limitations of the study

We identified three primary limitations of our strategy that should be considered when interpreting the results. First, the approaches that we have used to label ‘starter cell’ populations in the cerebellar cortex - we labeled on thee order of 1 x 1 mm of cerebellar cortical surface - are still coarse relative to the micro-zonal organization of the cerebellar cortex, which exhibits transitions at a spatial scale of 100-200 µm (Kostadinov et al., 2019; Oscarsson, 1979; Ozden et al., 2009; Schultz et al., 2009). Thus, the diversity of input and projection patterns that we observe within each region may be due to the heterogeneity of this starter population. Finer microzone-restricted labelling approaches (beyond the scope of this study) may reveal more precise targeting motifs. Second, our analysis of disynaptic connectivity was based on data with different signal-to-noise-ratio: disynaptic retrograde tracing labelled somas, which are easily counted, while disynaptic anterograde tracing labels axons, which are inherently harder to detect. Lower SNR may artificially hide structure and selectivity in the connections assayed. Third, our characterization of polysynaptic circuits is similarly constrained by an inability to label polysynaptic connections that are more than two synapses from the cerebellar cortex in the forebrain and by a lack of connectivity information with the spinal cord. Thus, we are not able to fully describe the long-range loops that the cerebellum may form with neocortex (Kelly & Strick, 2003), basal ganglia (Bostan & Strick, 2018), and spinal circuits (Sathyamurthy et al., 2020; Thanawalla et al., 2025). However, our approach allowed us, crucially, to unequivocally identify the intermediary relay hubs between disynaptic partner regions. Despite these limitations, our study provides a systematic interrogation of the brain-wide connectivity of cerebellar lobules V and simplex, two cerebellar regions with complementary preferences to encode action and reward. This work will serve as the starting point for interrogating how these cerebellar regions contribute to brain-wide computations for motor control and cognition. More generally, we describe a platform that can be utilized to delineate the mono- and disynaptic connectivity of any brain region, providing a general-purpose tool to relate local circuits to their long-range partners.

## Code and data availability

The *Braintracer* code, togerther with downsampled versions of the data, are availble on Github (https://github.com/samclothier/braintracer). *Braintracer* is also available for command-line installation as a public Python package on *Pypi* (https://pypi.org/project/braintracer/).

## Acknowledgements

We thank Soyon Chun, Michael Krumin, Caroline Reuter, Magda Robacha and Adam Tyson for technical support. This work was supported by grants from the Wellcome Trust (SHWF 221674 to L.F.R, *CDA 225951/Z/22/Z to D.K., PRF 201225/Z/16/Z and 224668/Z/21/Z to M.H.), the European Research Council (ERC AdG 695709 to M.H.)* by the Armenise-Harvard Foundation (CDA to L.F.R.), the Human Technopole (HT–ECF 3588 to L.F.R.).

## Author Contributions

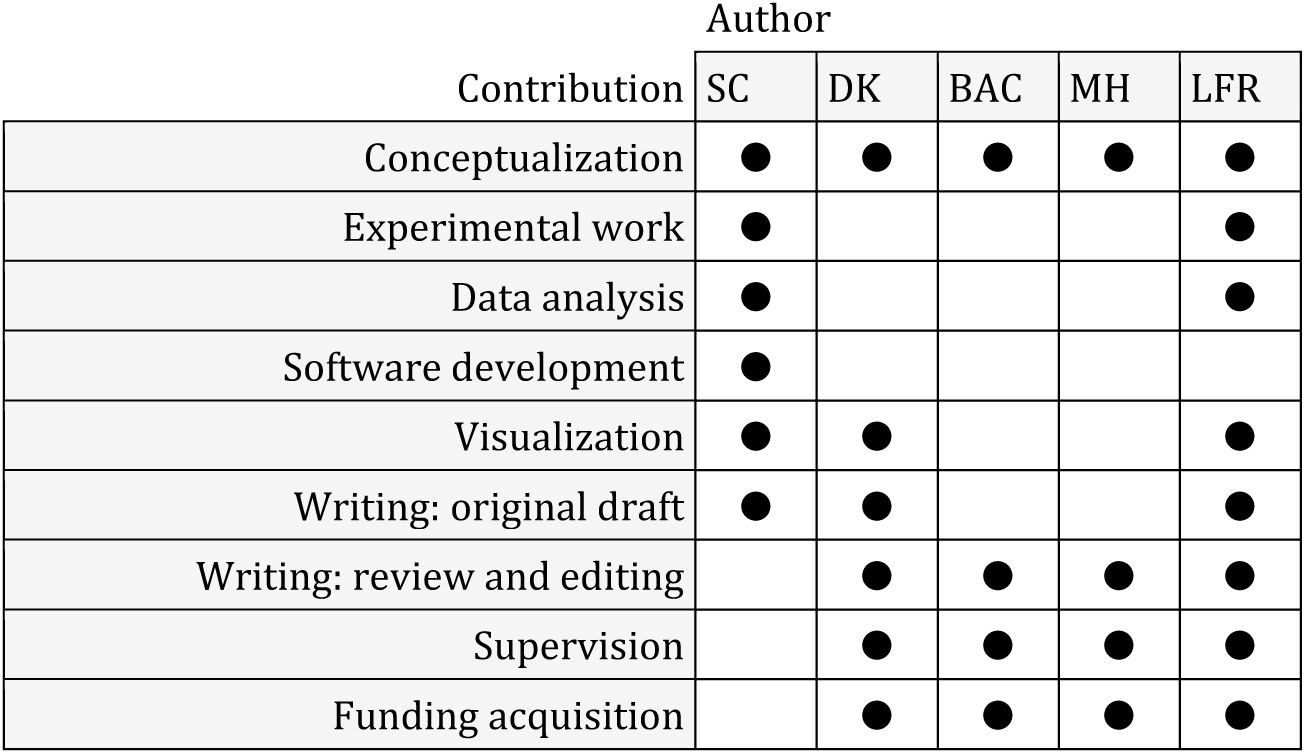

## Methods

All procedures were performed under and in accordance with the UK Animals Scientific Procedures Act (1986) under project and personal licenses issued by the Home Office following ethical review.

### Animals and Viral Vectors

Experiments were performed on 6-10 week-old mice of both sexes, maintained on a 12 hour light-dark cycle, at 20-24 °C and 45-65% humidity, in individually ventilated cages with food and water *ad libitum*. Twenty-one C57Bl/6J wild-type mice were used for TRIO experiments (https://www.jax.org/strain/013636). Eight Cre-dependent tdTomato reporter mice (Line Ai9, https://www.jax.org/strain/007909), expressing tdTomato conditional on the delivery of Cre recombinase, were used for anterograde tracing experiments. Three DAT-iCre mice (https://www.jax.org/strain/016583), expressing Cre in dopaminergic neurons, were used for experiments tracing the outputs of midbrain dopamine neurons; two Pcp2-Cre (https://www.jax.org/strain/010536) were used as controls for these experiments. The full list of viral vectors used in each experiment type is shown in **Table S2**.

### Surgical Procedures

Mice were anaesthetized with isoflurane (1–2% in oxygen), their body temperature was monitored and kept at 37–38 °C using a closed-loop heating pad, and the eyes were protected with ophthalmic gel (Viscotears Liquid Gel, Alcon). An analgesic (Rimadyl, 5 mg/kg) was administered subcutaneously before the procedure, and orally on subsequent days. Whenever the procedure exposed the brain, dexamethasone (0.5 mg/kg, IM) was administered intramuscularly 30 min before the procedure to prevent brain oedema. The exposed brain was constantly perfused with artificial cerebrospinal fluid (150 mM NaCl, 2.5 mM KCl, 10 mM HEPES, 2 mM CaCl2, 1 mM MgCl2; pH 7.3 adjusted with NaOH, 300 mOsm). The head was shaved and disinfected; the cranium was exposed via a scalp incision and leveled in the DV and ML axes; small craniotomies were opened with a dental drill.

Unilateral stereotaxic injections (coordinates in **Table S1**) were delivered through the small craniotomies (Φ = 0.5 mm) with a beveled glass micropipette (Φ = ∼20 µm tip) at a rate of 20 nl/min with manual syringe pressure (volumes described below). Micropipettes were left in position for approximately 5 minutes before being retracted. Craniotomies were protected with mineral oil before suturing the scalp with veterinary glue (Vetbond, 3M).

### Retrograde transsynaptic tracing

Retrograde transsynaptic tracing of the monosynaptic inputs to olivary neurons projecting to specific cerebellar cortical lobules was achieved with the TRIO method (Schwarz et al., 2015). 100 nl of a Cre-expressing retrograde virus, either canine adenovirus-2 (CAV2)-CMV-Cre (1x10^12^ GC/mL) (Kremer et al., 2000) or AAVrg-hSyn-Cre (7x10^12^ GC/mL, Addgene 105553) (Lavoie & Liu, 2020; Tervo et al., 2016) was injected into the target cerebellar lobule (LS, LV) to infect incoming climbing fibers and retrogradely drive Cre expression in their parent IO neurons.. In a subset of these experiments (N=8), to visualise and validate the targeting and spread of the injection, we co-injected a constitutive virus expressing a blue fluorescent protein (AAV2/1-EF1a-EBFP2, 2*10^12^ GC/mL) to express EBFP2 in local neurons. In the same surgery, we then performed a second 200 nl injection in the IO of the Cre-dependent molecular toolkit for transsynaptic retrograde rabies tracing: AAV2/1-EF1a-(ATG-out)-FLEX-EGFP (3*10^12^ GC/mL) mixed with either AAV2/1-Syn-FLEX-TVA-2A-oG (6*10^12^ GC/mL) or AAV2/1-EF1a(ATG-out)-FLEX-TVA-P2A-N2cG (6*10^12^ GC/mL). This mix primed the olivary neurons for infection with a modified rabies virus, conditional on their retrograde transduction with Cre, and labeled them with EGFP. (**Figure 5e-g**). After 3 weeks, we re-targeted the same site in the IO with a 300 nl injection of an engineered rabies virus, either EnvA-dG-SADB19RV-mCherry (1x10^9^ CFU/mL) or EnvA-dG-N2cRV-tdTomato (1x10^8^ CFU/mL), which infected the competent IO neurons and drove expression of a red fluorescent protein in receptive IO neurons and in thousands of their presynaptic inputs (Wickersham et al., 2007). For maximal yield (Rossi et al., 2020), the animals were then sacrificed 1 and 2 weeks following rabies injection for SADB19 or N2C variants.

### Anterograde transsynaptic tracing

Anterograde mono- and disynaptic tracing of the outputs of neurons in the cerebellar cortex was achieved with injections of the anterograde AAV2/1-hSyn-Cre (100 nL, 10^13^ GC/ml) into the target cerebellar cortical lobule (either LS or LV) in Ai9 reporter mice, where delivery of Cre drives the expression of the red fluorescent protein tdTomato. Aside from labelling most neurons at the injection site and their axons, this virus spreads transsynaptically (Zingg et al 2017; 2020) also labelling the postsynaptic neurons in the cerebellar nuclei and their axonal projections. In some experiments, to visualize and validate the targeting and spread of the injection, we co-injected a constitutive virus (AAV2/1-EF1a-EBFP2) expressing blue fluorescent protein in local neurons.

Tracing of dopaminergic neurons from the VTA to the IO was achieved with an injection of AAV2/1-FLEX-tdTomato virus into the VTA of DAT-iCre mice. Animals were sacrificed for imaging 3 weeks after injection.

### Block-face 2P tomography

Fluorescent brains were imaged with block-face two-photon tomography to map labelled neurons. Animals were deeply anesthetized with ketamine-xylazine and perfused transcardially with PBS followed by 4% paraformaldehyde in PBS (PB, pH 7.4). The brain was extracted and post-fixed in 4% PFA-0.1 M PB for 24 hours at 10°C. Fixed brains were stored in PBS prior to imaging.

Whole brain 3D imaging was performed using a custom-built serial two-photon tomography microscope coupled to a microtome (Ragan et al., 2012). The fixed brain was aligned on a mold and mounted in 0.5% agar-PBS; then, it was glued on the microtome base on the frontal side. The 40 µm thick coronal volumes were serially imaged at either 980 or 780 nm (50-100 mW) with an X-Y pixel size of 2 µm, collecting optical slices every 5 µm (or 2 µm for VTA axons in the IO); 40 µm thick physical sections were then sliced off before resuming imaging. The acquisition was controlled with BakingTray and the resulting tiled image stacks were pre-processed with Stitchit (https://github.com/SWC-Advanced-Microscopy).

### Analysis and quantification with Braintracer

3D imaging stacks were processed with our new custom analysis pipeline, *Braintracer* (https://github.com/samclothier/braintracer). Braintracer integrates with and expands on packages provided by the *BrainGlobe* API (Claudi et al., 2020), including 3D registration to atlases (BrainReg), automatic cell detection (Cellfinder), and provides two workflows to get from imaging to data visualization and quantification: automatic neuron detection and brain-wide fluorescence analysis (**Supplementary Figure 1**).

### Detection of Brain-wide neurons

First, brain volumes were registered with a 3D transform to the Allen Common Reference atlas, using *Brainreg* (Niedworok et al., 2016) (**Supplementary Figure 1a-b**).

Second, brain-wide traced neurons were automatically detected and classified with *Cellfinder* (**Supplementary Figure 1**) (Tyson et al., 2021). Since the standard convolutional neural network (50-layer ResNet) yielded excessive false-positives, we optimized its performance by re-training the classifier on ∼1,000 manually validated neurons from our own datasets, including examples from multiple brain areas, thus increasing accuracy while maintaining generality. This workflow was able to automatically detect and classify neurons of vastly different morphology across brain regions (**Supplementary Figure 5**), with few false positives and negatives (validation accuracy of 0.92 and a loss of 0.19). Notably, our custom-trained CNN successfully rejected bright artefacts at the surface of the brain and in the ventricles. The only exception were IO neurons in a few TRIO experiments in which only the red and green channels had been imaged; since cell detection relies on the comparison with an autofluorescence channel to discard noise, classification performs poorly when a cell has emission in both channels. We therefore performed manual counts in the IO for datasets where only the red and green channels were acquired. Finally, the coordinates of the identified cells were mapped in the Allen Common Reference Atlas coordinates using the 3D transform computed on the imaging volume (**Supplementary Figure 1**).

For 2D visualizations, whole brain tracing counts were discretized in 50x50x50 µm voxels, then smoothed with a 3D Gaussian kernel (sigma = 0.5), normalized to yield a connection probability map, and finally averaged along the projection axis of interest. Summary maps represent pixel-wise medians across experiments.

### Automatic Detection of Brain-wide Fluorescence Signal

The analysis of anterograde projection datasets relied on measuring fluorescence of axonal projections (Oh et al, 2014). After 3D brain registration to the Allen Common Reference Atlas, images were cleaned by a brain-wide subtraction of autofluorescence (**Supplementary Figure 1**). This involved fitting a linear RANSAC (RANdom Sample Consensus) regression model to the relationship between the signal channel (red) and background channel (blue, or green when blue not imaged). Since autofluorescence produces correlated signals in all channels, while experimental labelling was sparse, this model (*y=mx+c*) captured varying autofluorescence across brain areas (mx) as well as the baseline brightness of the sample (+c). We then obtained a true signal value for each pixel by subtracting the fitted autofluorescence component:

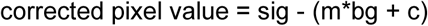

For each slice, we considered true labelling pixels with signal above the 99th percentile, and then applied a median filter (3*3) to emphasize axon-sized components over thresholded noise. The coordinates of labelled pixels were then mapped in the Allen Common Reference Atlas coordinates using the 3D transform computed on the imaging volume and quantified in terms of voxel volume in mm³ (**Supplementary Figure 1i**).

For 2D visualizations, whole brain tracing counts were discretized in 50x50x50 µm voxels, then smoothed with a 3D Gaussian kernel (sigma = 0.5), normalized to yield a connection probability map, and finally averaged along the projection axis of interest. Summary maps were pixel-wise medians across experiments.

### Additional subdivision for the Inferior Olive and Mesodiencephalic Junction

The inferior olive and the mesodiencephalic junction (MDJ) are two key regions for cerebellar circuits, but their anatomical divisions were previously missing from the Allen Common Reference Atlas. Thus, we augmented the Allen Common Reference Atlas with new annotations for the subdivisions of the inferior olive (based on Luo et al. (2023)) and a region for the MDJ (based on Wang et al. (2022)) that was previously classified as midbrain. This augmented brain atlas is provided for use with Braintracer (**Supplementary Figure 2**).

### Region Selection

Tracing counts (cell number or voxel volume) were quantified for each brain subdivision in the Allen Common Reference Atlas, where areas are ordered according to a hierarchical tree. For clarity, we included in our analysis only areas containing a substantial fraction of the total signal. Then, to select the appropriate granularity of analysis for the major brain domains - brainstem, pons, cerebellum, midbrain, forebrain - we iteratively split parent areas into child branches and stopped when further splitting would reduce the number of inputs below 0.5% of the total number of inputs as a mean across all datasets.

For monosynaptic MF inputs to the cerebellum, we restricted our search to brain areas known to receive projections from the DCN, according to the Allen Common Reference atlas. These included regions in the pons, midbrain, and medulla, including the trigeminal nuclei, tegmental reticular nucleus and vestibular nuclei. For the disynaptic output analysis, to determine the granularity for output regions we relaxed the threshold for subregion selection to 0.01% of total labelling and considered the following regions: thalamus, superior colliculus sensory and motor parts, red nucleus, periaqueductal gray, and VTA.

### Statistics, Connection and spatial selectivity index

To verify whether connectivity differed as a function of experimental group — LV versus LS — anatomical area, and the interaction between group and area, we used a two-way ANOVA, which asks whether the LV–LS difference depends on anatomical location. To reveal differences between areas across groups, we followed up significant ANOVA group or interaction effects with post-hoc pairwise U-tests.

To compare connectivity patterns across experimental groups, we computed two metrics: a connection selectivity index and a topographic selectivity index. For each area *a*, the connection selectivity index (rS) quantified differences in connection strength (C) across the two experimental groups:

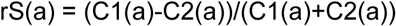

rS is bounded between [-1, 1]; rS close to -1 or 1 corresponds to dominant connectivity from one experimental group; rS close to 0 corresponds to similar connection strengths across groups.

The spatial selectivity index (tS) instead was a proxy of the similarity between the spatial distribution of labelling within a given region between experimental groups. For each area, it was measured as the fraction of 3D voxels that contained at least 10 times as much labelling in one group than the other.

---

**Supplementary Figure 1:**
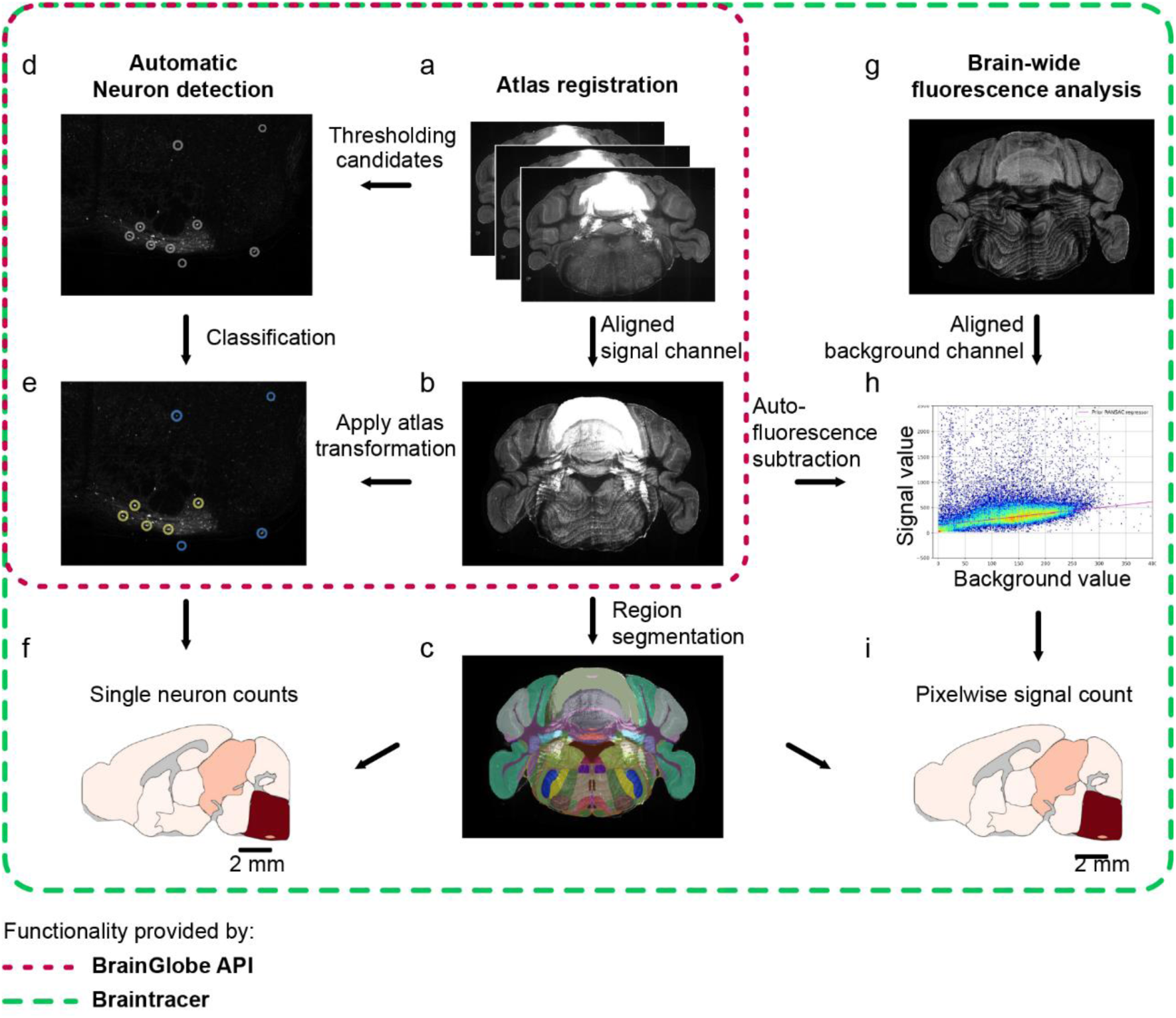
A custom high-throughput processing pipeline for analysis of high-resolution serial section two-photon imaging datasets. (**a**) Raw image stack obtained from serial 2-photon tomography. (**b**) Dataset registered to the Allen Common Reference atlas in 3D. (**c**) Atlas regions overlaid on the registered data. (**d**) Cellfinder applies a threshold to identify cell candidates. (**e**) Cells are classified with a deep neural network. (**f**) Identified cells are transformed into atlas space using the transformation from b and assigned to brain regions for analysis of brain-wide cerebellar inputs and outputs. (**g**) Registered background channel is compared to the foreground channel to identify autofluorescence. (**h**) Autofluorescence is removed from the registered data in **b** by computing the pixel-wise subtraction of the average difference in brightness between the signal and background (reference) channels. This difference is found by fitting a robust regression model to a sample of pixels in the signal and background channels. Using this fit, if a pixel position has a high value in both channels (i.e. is autofluorescent), it will be subtracted from the image. (**i**) Following subtraction, a threshold is applied and the mean per-region fluorescence is calculated for analysis of brain-wide cerebellar inputs and outputs.

**Supplementary Figure 2:**
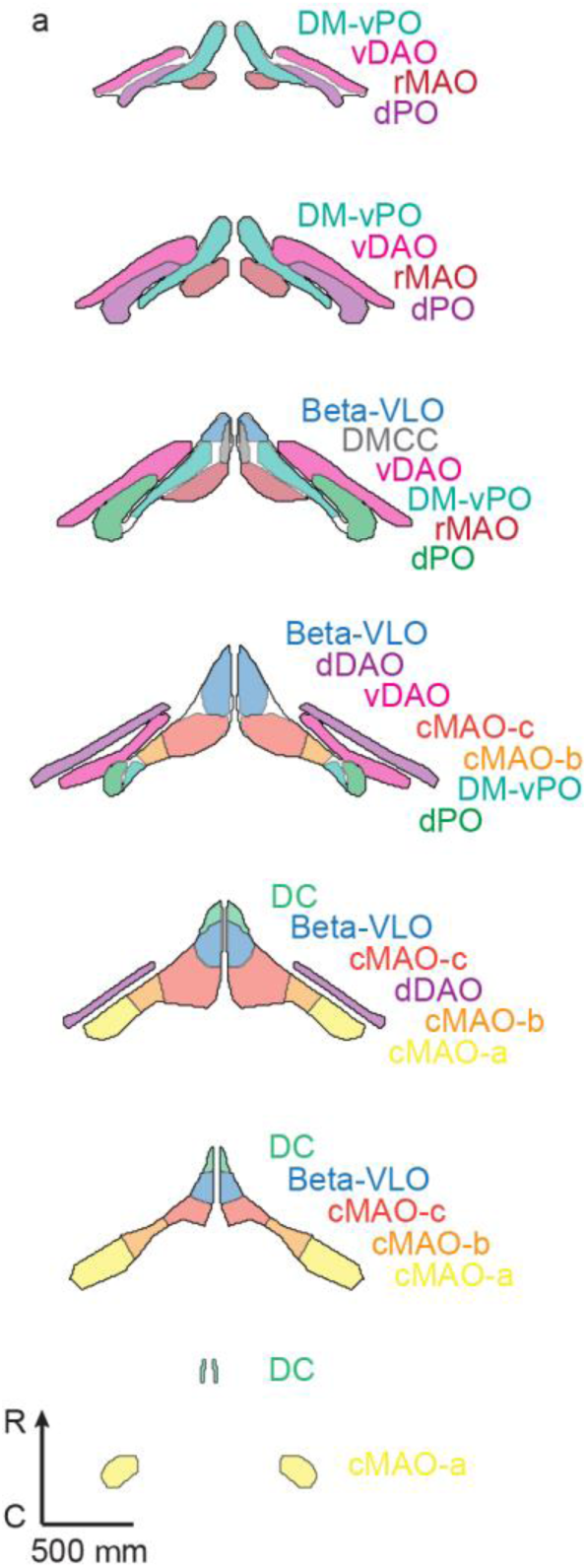
New atlas annotations of inferior olive subnuclei. (**a**) Rostral (top) to caudal (bottom) slices through the Braintracer augmented Allen Common Reference atlas, demonstrating the new annotations for inferior olive subnuclei: dorsomedial subnucleus - ventral lamella of the principal olive (DM-vPO, rostral medial accessory olive (rMAO), dorsal principal olive (dPO), subnucleus beta - ventrolateral outgrowth subnucleus (Beta-VLO), Dorsomedial cell column subnucleus (DMCC), ventral fold of the dorsal accessory olive (vDAO), dorsal fold of the dorsal accessory olive (dDAO), subnuclei a, b, and c of the caudal part of the medial accessory olive (cMAO-a, -b, and -c), dorsal cap subnucleus (DC).

**Supplementary Figure 3:**
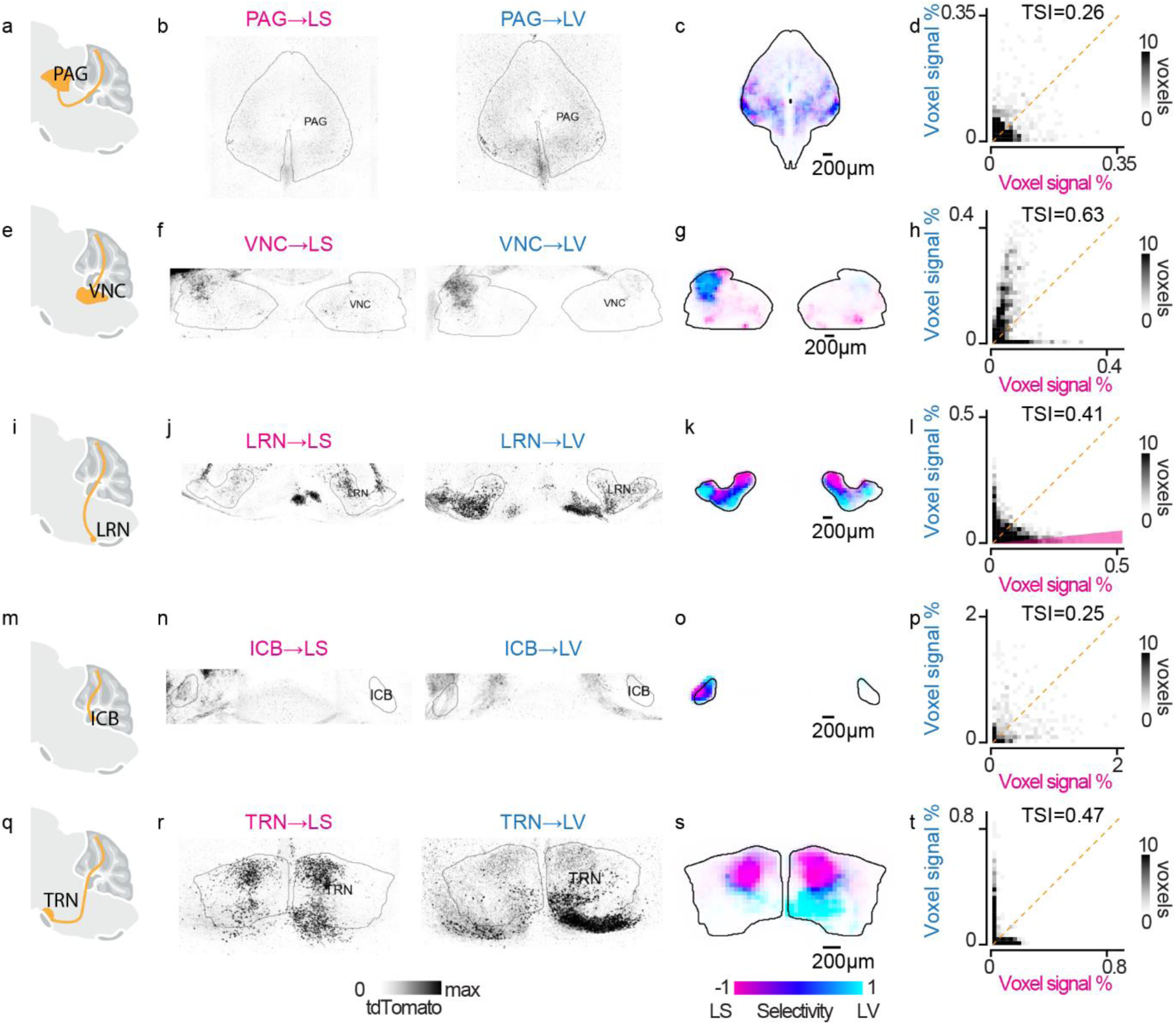
Additional monosynaptic input regions. (**a**) Schematic of anterograde labeling of monosynaptic MF inputs from the PAG. (**b**) Example maximum projections of MF inputs in the PAG for brains injected in LS (left) and LV (right). (**c**) Coronal projection of the average labeling probability density (voxel-wise median) in the PAG for LS (n=4, magenta) and LV (n=4, cyan) injections, mapped into HSV color space where LS-exclusive voxels are magenta, LV-exclusive are cyan, and 50/50 mixed voxels are blue. Data from each experiment were aligned to the Allen Common Reference atlas before averaging. (**d**) Normalized labelling density of voxels in the PAG from experiments targeting LV (cyan, y axis) plotted against matching voxels from injections in LS (magenta, x axis). (**e**, **f**, **g**, **h**) Same as **a**-**d**, for MFs in VNC. (**i**, **j**, **k**, **l**) Same as **a**-**d**, for MFs in LRN. (**m**, **n**, **o**, **p**) Same as **a**-**d**, for MFs in ICB. (**q**, **r**, **s**, **t**) Same as **a**-**d**, for MFs in TRN.

**Supplementary Figure 4:**
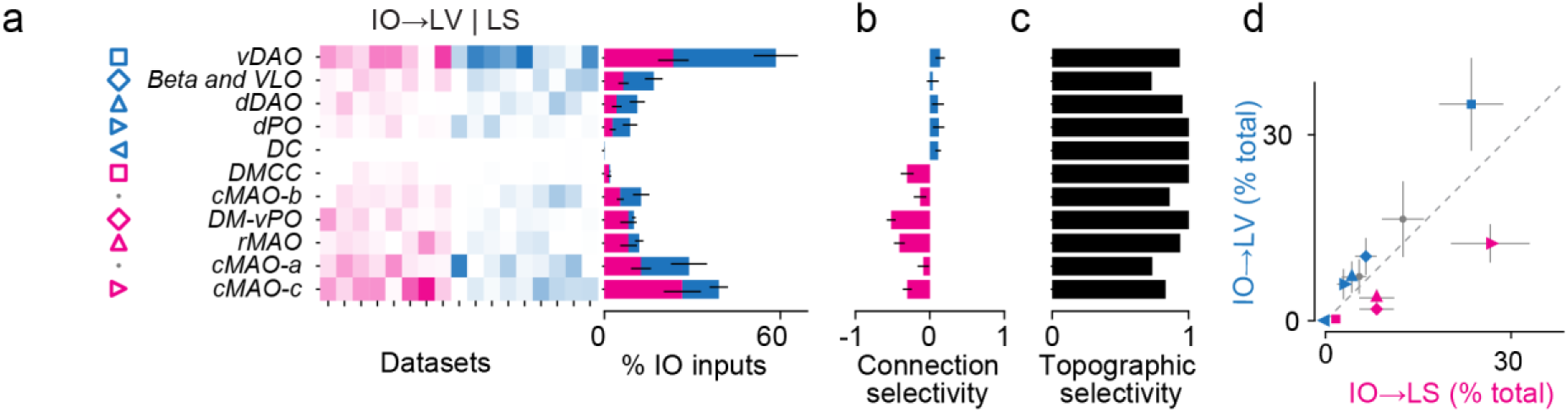
TRIO starter neurons and local inputs from the IO. (**a**) Distribution of LV- and LS-projecting CFs across olivary subdivisions, and other disynaptic inputs from the IO. Left: Normalized inferior olive neurons labeled in LS (magenta) or LV (cyan) from all IO subdivisions (rows) in each dataset (column, sorted by experimental groups). Right: stacked histogram of average normalized inferior olive neuron starter cells across experimental groups (mean ± s.e.m. is shown for each group). (**b**) Average connection selectivity index of the input strength from each area (mean ± s.e.m. for all pairwise comparisons between datasets across experimental groups). Negative values indicate stronger labelling in LS datasets (magenta); positive values indicate stronger labelling in LV datasets (cyan). (**c**) Spatial selectivity index for the labelling density in each area. Higher values correspond to larger differences in the spatial distribution of labelling. (**d**) Average (mean ± s.e.m.) labelling intensity in LS vs LV experimental groups for olivary subdivisions shown in **a**-**c**, where areas with significant labeling differences have colored markers.

**Supplementary Figure 5:**
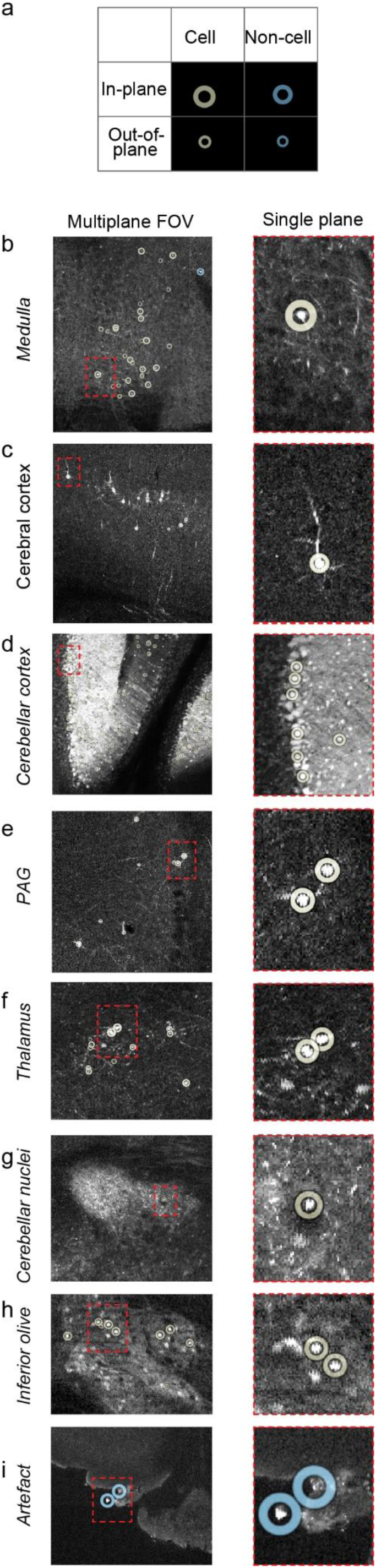
Extracting soma locations from serial section images using a deep neural network classifier. (**a**) Legend describing markers for cell candidates classified as ‘cell’ and ‘non-cell’. Out-of-plane markers are smaller to indicate distance in z from the focal plane. (**b**) Cell classifications from an example dataset injected with the retrograde TRIO approach in the medulla. Left: Whole FOV including cells up to 5 planes away. Right: Inset view only including classifications on the focal plane. (**c**) Same as **b**, for cells in the cerebral cortex. (**d**) Same as **b**, for cells in the cerebellar cortex (Cbx). (**e**) Same as **b**, for cells in the periaqueductal grey (PAG). (**f**) Same as **b**, for cells in the thalamus (TH). (**g**) Same as **b**, for cells in the deep cerebellar nuclei (CbN). (**h**) Same as **b**, for cells in the inferior olive. (**i**) Same as **b**, for autofluorescence artefacts along the surface of the sample. These are correctly rejected as non-cells by the classifier.

**Supplementary Figure 6:**
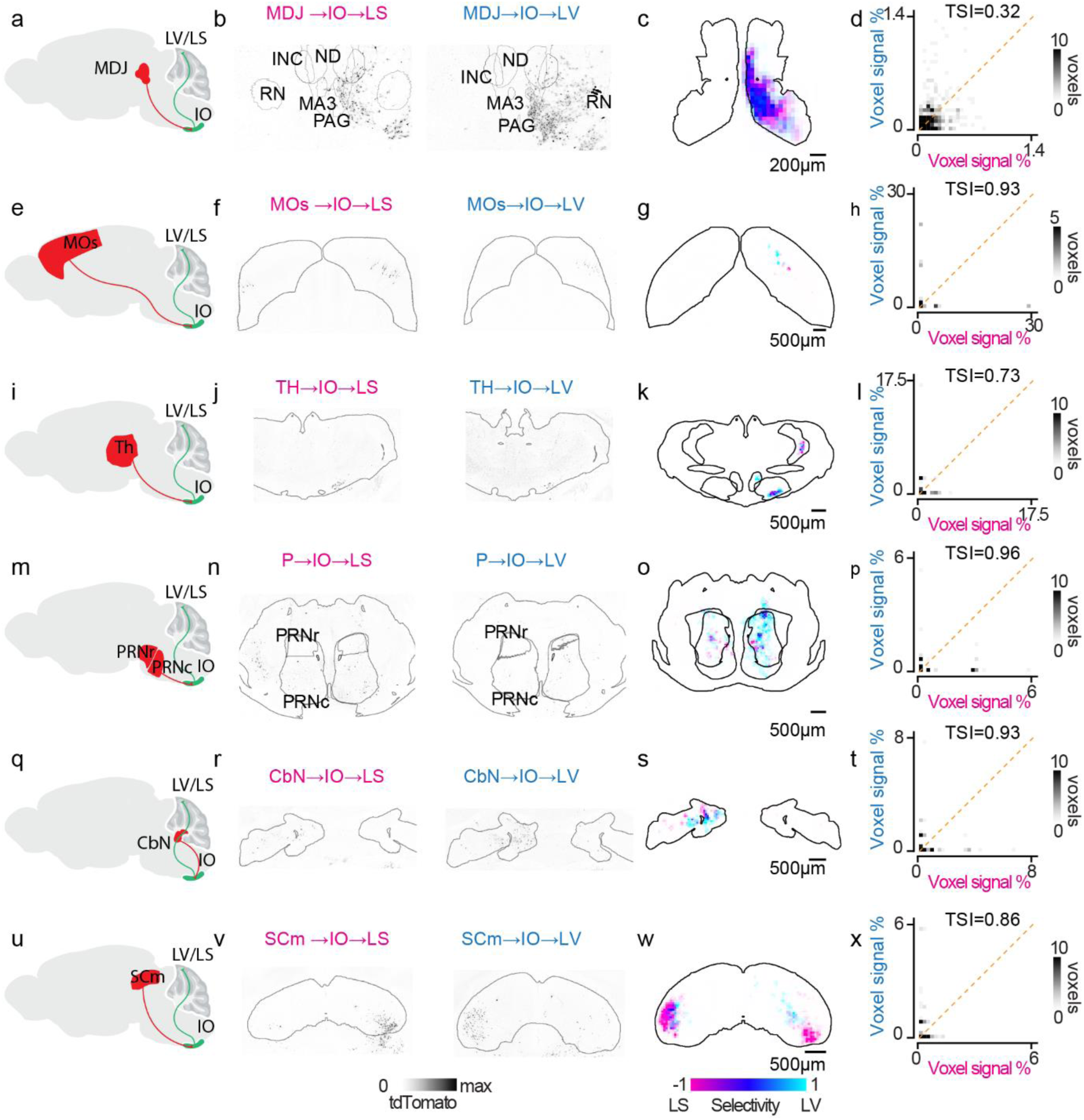
Additional disynaptic input regions. (**a**) Schematic of TRIO retrograde labeling of disynaptic inputs to LS and LV via the IO. (**b**) Example maximum projections of disynaptic inputs in the MDJ for brains injected in LS (left) and LV (right). (**c**) Coronal projection of the average labeling probability density (voxel-wise median) in the MDJ for LS (n=8, magenta) and LV (n=9, cyan) injections, mapped into HSV color space where LS-exclusive voxels are magenta, LV-exclusive are cyan, and 50/50 mixed voxels are blue. Data from each experiment were aligned to the Allen Common Reference atlas before averaging. (**d**) Normalized labelling density of voxels in the MDJ from experiments targeting LV (cyan, y axis) plotted against matching voxels from injections in LS (magenta, x axis). (**e**, **f**, **g**, **h**) Same as **a**-**d**, for disynaptic inputs in MOs. (**i**, **j**, **k**, **l**) Same as **a**-**d**, for disynaptic inputs in TH. (**m**, **n**, **o**, **p**) Same as **a**-**d**, for disynaptic inputs in P. (**q**, **r**, **s**, **t**) Same as **a**-**d**, for disynaptic inputs in CbN. (**u**, **v**, **w**, **x**) Same as **a**-**d**, for disynaptic inputs in SCm.

**Supplementary Figure 7:**
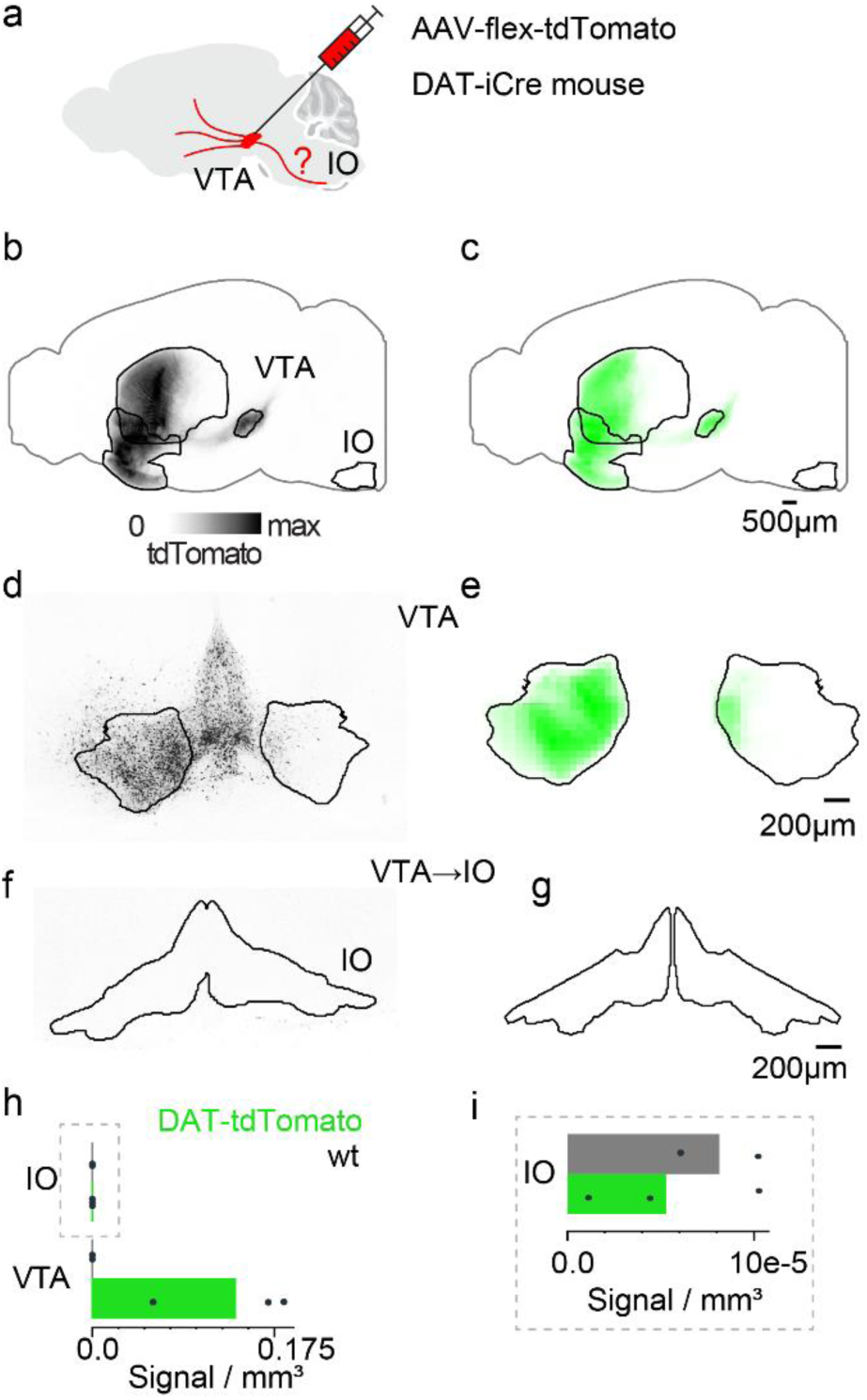
Lack of dopaminergic axons in the inferior olive. (**a**) Schematic AAV2/1-FLEX-tdTomato injection into the VTA, expressing tdTomato in dopaminergic axons. (**b**) Background subtracted sagittal projection of example brain with labelled dopaminergic projections after VTA injection. (**c**) Sagittal projection of the median labelling density of injected DAT-iCre animals (n=3). Each brain was aligned to the Allen Common Reference atlas before averaging. (**d**) Example coronal slice through the VTA with labelled dopaminergic neurons. (**e**) Same as **c**, showing labelling density within the VTA. (**f**) Example coronal slice through the IO, lacking axonal labelling from VTA dopaminergic projections. (**g**) Same as **c**, showing labelling density within the IO. (**h**) Mean labeling volume (bar plots), and individual datasets (points), in the inferior olive (IO) and ventral tegmental area (VTA), for animals injected with AAV-flex-tdTomato (green, n=3) or control animals (gray, n=2). (**i**) Inset from **h**, magnifying labelling counts for the IO.

**Table S1.** Stereotaxic coordinates of injection sites. Coordinates are measured from Lambda. AP: anteroposterior axis, ML: mediolateral axis.

| Injection site | AP (mm) | ML (mm) | Depth (mm) |
| --- | --- | --- | --- |
| Lobule Simplex | -2.30 | -2.30 | 0.25 |
| Lobule V | -2.00 | -0.80 | 0.15 |
| *Inferior olivary*<br>*complex (contralateral)* | *-2.80* | *0.40* | *5.40* |

**Table S2.** Viruses used across all procedures and experiments. Stock titer and injection titers given as GC/ml (genome copies) or IU/ml (infective units). Each item was obtained from the indicated origin institution.

| Virus | Titer (stock) | Titer (injection) | Origin |
| --- | --- | --- | --- |
| AAVretro-Cre | $\sim 7 \times 10^{12}$ GC/ml | $\sim 7 \times 10^{12}$ GC/ml | Addgene (ID: 24593) |
| CAV-Cre | $\sim 1 \times 10^{12}$ GC/ml | $\sim 1 \times 10^{12}$ GC/ml | PVM (Plateau IGMM) |
| AAV2/1-EF1a-(ATG-out)-<br>FLEX-EGFP | $\sim 1 \times 10^{14}$ GC/ml | $\sim 1 \times 10^{12}$ GC/ml | SWC Vector Core |
| AAV2/1-Syn-FLEX-TVA-2A-oG | $\sim 8 \times 10^{12}$ GC/ml | $\sim 6 \times 10^{12}$ GC/ml | Salk Institute Vector<br>Core |
| AAV2/1-EF1a-(ATG-out)-<br>FLEX-TVA-P2A-N2cG | $\sim 1 \times 10^{14}$ GC/ml | $\sim 6 \times 10^{12}$ GC/ml | SWC Vector Core |
| EnvA-dG-SADB19RV-mCherry | $\sim 1 \times 10^{11}$ IU/ml | $\sim 1 \times 10^{11}$ IU/ml | NTNU Vector Core |
| EnvA-dG-N2cRV-tdTomato | $\sim 1 \times 10^7$ IU/ml | $\sim 1 \times 10^7$ IU/ml | NTNU Vector Core |
| AAV2/1-EF1a-EBFP2 | $\sim 1 \times 10^{14}$ GC/ml | $\sim 1 \times 10^{12}$ GC/ml | SWC Vector Core |
| AAV2/1-hSyn-Cre | $\sim 2 \times 10^{13}$ GC/ml | $\sim 1 \times 10^{13}$ GC/ml | Addgene (ID: 105553) |

**Table S3.** Other reagents and equipment. Reagents for perfusion, injectable drugs for stereotaxic surgeries, and notable equipment used. Stock and injectable concentrations indicated. Each item was obtained from the indicated origin company.

| Item | Concentration (stock) | Concentration (injection) | Origin |
| --- | --- | --- | --- |
| Paraformaldehyde | 16% | 4% |  |
| Phosphate-buffered saline | 10x | 1x | Sigma-Aldrich Chemie GmbH |
| Liquid isoflurane | 100% w/w | 2% (inhalation) | Zoetis UK Limited |
| Dexamethasone | 2 mg/ml | 2 mg/ml | MSD Animal Health |
| Carprofen | 50 mg/ml | 5 mg/kg | Zoetis UK Limited |
| Ketamine-xylazine mix | 80-6 mg/ml | 80-6 mg/ml | Sigma-Aldrich Chemie GmbH |
| Saline | 9 mg/ml NaCl | 9 mg/ml | Animalcare Ltd |
| Blaubrand IntraMARK Glass micropipettes |  |  | Brand GmbH |
| Stereotaxic frame |  |  | Narishige Scientific Instrument Lab |
| Eye ointment |  |  | Bayer Vital GmbH |
| Dermafuse |  |  | Millpledge Veterinary |

## References

1. Apps, R., & Garwicz, M. (2005). Anatomical and physiological foundations of cerebellar information processing. Nat Rev Neurosci, 6(4), 297–311. 10.1038/nrn1646

2. Apps, R., & Hawkes, R. (2009). Cerebellar cortical organization: a one-map hypothesis. Nat Rev Neurosci, 10(9), 670–681. 10.1038/nrn2698

3. Apps, R., Hawkes, R., Aoki, S., Bengtsson, F., Brown, A. M., Chen, G., Ebner, T. J., Isope, P., Jorntell, H., Lackey, E. P., Lawrenson, C., Lumb, B., Schonewille, M., Sillitoe, R. V., Spaeth, L., Sugihara, I., Valera, A., Voogd, J., Wylie, D. R., & Ruigrok, T. J. H. (2018). Cerebellar Modules and Their Role as Operational Cerebellar Processing Units: A Consensus paper [corrected]. Cerebellum, 17(5), 654–682. 10.1007/s12311-018-0952-3

4. Bostan, A. C., & Strick, P. L. (2018). The basal ganglia and the cerebellum: nodes in an integrated network. Nat Rev Neurosci, 19(6), 338–350. 10.1038/s41583-018-0002-7

5. Claudi, F., Petrucco, L., Tyson, A. L., Margrie, T. W., Portugues, R., & Branco, T. (2020). BrainGlobe Atlas API: A common interface for neuroanatomical atlases. Journal of Open Source Software, 5(54), 2668. 10.21105/joss.02668

6. Dacre, J., Colligan, M., Clarke, T., Ammer, J. J., Schiemann, J., Chamosa-Pino, V., Claudi, F., Margrie, T. W., & Duguid, I. (2021). A cerebellar-thalamocortical pathway drives behavioral context-dependent movement initiation. Neuron, 109(14), 2302–2314.e2308. 10.1016/j.neuron.2021.05.018

7. Eccles, J. C., Ito, M., & Szentágothai, J. (1967). The cerebellum as a neuronal machine. Springer Science & Business Media.

8. Fallon, J. H., & Bañales, J. L. (1984). Neurons in the ventral tegmentum have separate populations projecting to telencephalon and inferior olive, and to superior colliculus and lower brain stem in the rat. Brain Research, 321(2), 332–336. https://www.sciencedirect.com/science/article/abs/pii/0006899384901884?via%3Dihub

9. Finkelstein, A., Daie, K., Rozsa, M., Darshan, R., & Svoboda, K. (2026). Connectivity underlying motor cortex activity during goal-directed behaviour. Nature, 649(8096), 416–422. 10.1038/s41586-025-09758-6

10. Fujita, H., Kodama, T., & du Lac, S. (2020). Modular output circuits of the fastigial nucleus for diverse motor and nonmotor functions of the cerebellar vermis. Elife, 9. 10.7554/eLife.58613

11. Gao, Z., Davis, C., Thomas, A. M., Economo, M. N., Abrego, A. M., Svoboda, K., De Zeeuw, C. I., & Li, N. (2018). A cortico-cerebellar loop for motor planning. Nature, 563(7729), 113–116. 10.1038/s41586-018-0633-x

12. Heffley, W., & Hull, C. (2019). Classical conditioning drives learned reward prediction signals in climbing fibers across the lateral cerebellum. Elife, 8, e46764. 10.7554/eLife.46764

13. Heffley, W., Song, E. Y., Xu, Z., Taylor, B. N., Hughes, M. A., McKinney, A., Joshua, M., & Hull, C. (2018). Coordinated cerebellar climbing fiber activity signals learned sensorimotor predictions. Nat Neurosci, 21(10), 1431–1441. 10.1038/s41593-018-0228-8

14. Henschke, J. U., & Pakan, J. M. P. (2020). Disynaptic cerebrocerebellar pathways originating from multiple functionally distinct cortical areas. Elife, 9, e59148. 10.7554/eLife.59148

15. Herzfeld, D. J., & Lisberger, S. G. (2025). Neural circuit mechanisms to transform cerebellar population dynamics for motor control in monkeys. bioRxiv. 10.1101/2025.02.21.639459

16. Huang, C. C., Sugino, K., Shima, Y., Guo, C., Bai, S., Mensh, B. D., Nelson, S. B., & Hantman, A. W. (2013). Convergence of pontine and proprioceptive streams onto multimodal cerebellar granule cells. Elife, 2, e00400. 10.7554/eLife.00400

17. Ito, M. (2008). Control of mental activities by internal models in the cerebellum. Nat Rev Neurosci, 9(4), 304–313. 10.1038/nrn2332

18. Kelly, R. M., & Strick, P. L. (2003). Cerebellar loops with motor cortex and prefrontal cortex of a nonhuman primate. J Neurosci, 23(23), 8432–8444. 10.1523/JNEUROSCI.23-23-08432.2003

19. Kim, M. H., Znamenskiy, P., Iacaruso, M. F., & Mrsic-Flogel, T. D. (2018). Segregated Subnetworks of Intracortical Projection Neurons in Primary Visual Cortex. Neuron, 100(6), 1313–1321 e1316. 10.1016/j.neuron.2018.10.023

20. Kostadinov, D., Beau, M., Blanco-Pozo, M., & Hausser, M. (2019). Predictive and reactive reward signals conveyed by climbing fiber inputs to cerebellar Purkinje cells. Nat Neurosci, 22(6), 950–962. 10.1038/s41593-019-0381-8

21. Kostadinov, D., Clark, B. A., & Hausser, M. (2025). Fast and slow learning mediated by distinct climbing fiber signals. bioRxiv. 10.64898/2025.12.19.695643

22. Kostadinov, D., & Häusser, M. (2022). Reward signals in the cerebellum: Origins, targets, and functional implications. Neuron, 110(8), 1290–1303. 10.1016/j.neuron.2022.02.011

23. Kremer, E. J., Perricaudet, M., Yeh, P., Moullier, P., Chartier, C., Chillon, M., Curiel, D. T., Bergelson, J., Engelhardt, J. F., & Bramson, J. L. (2000). Canine adenovirus vectors: An alternative for adenovirus-mediated gene transfer. Journal of Virology, 74(1), 505–512. 10.1128/JVI.74.1.505-512.2000

24. Lavoie, A., & Liu, B. H. (2020). Canine Adenovirus 2: A Natural Choice for Brain Circuit Dissection. Front Mol Neurosci, 13, 9. 10.3389/fnmol.2020.00009

25. Lee, K. H., Mathews, P. J., Reeves, A. M. B., Choe, K. Y., Jami, S. A., Serrano, R. E., & Otis, T. S. (2015). Circuit mechanisms underlying motor memory formation in the cerebellum. Neuron, 86(2), 529–540. 10.1016/j.neuron.2015.03.010

26. Leergaard, T. B., & Bjaalie, J. G. (2007). Topography of the complete corticopontine projection: from experiments to principal Maps. Front Neurosci, 1(1), 211–223. 10.3389/neuro.01.1.1.016.2007

27. Li, N., & Mrsic-Flogel, T. D. (2020). Cortico-cerebellar interactions during goal-directed behavior. Curr Opin Neurobiol, 65, 27–37. 10.1016/j.conb.2020.08.010

28. Luo, Y., Chao, Y., Owusu-Mensah, R. N. A., Zhang, J., Hirata, T., & Sugihara, I. (2023). Neurogenic timing of the inferior olive subdivisions is related to the olivocerebellar projection topography. Sci Rep, 13(1), 7114. 10.1038/s41598-023-33497-1

29. Madisen, L., Zwingman, T. A., Sunkin, S. M., Oh, S. W., Zariwala, H. A., Gu, H., Ng, L. L., Palmiter, R. D., Hawrylycz, M. J., Jones, A. R., Lein, E. S., & Zeng, H. (2010). A robust and high-throughput Cre reporting and characterization system for the whole mouse brain. Nature Neuroscience, 13(1), 133–140. 10.1038/nn.2467

30. Niedworok, C. J., Brown, A. P. Y., Jorge Cardoso, M., Ourselin, S., Modat, M., & Margrie, T. W. (2016). aMAP is a validated pipeline for registration and segmentation of high-resolution mouse brain data. Nature Communications, 7, 11879. 10.1038/ncomms11879

31. Oh, S. W., Harris, J. A., Ng, L., Winslow, B., Cain, N., Mihalas, S., Wang, Q., Lau, C., Kuan, L., Henry, A. M., Mortrud, M. T., Ouellette, B., Nguyen, T. N., Sorensen, S. A., Slaughterbeck, C. R., Wakeman, W., Li, Y., Feng, D., Ho, A., Zeng, H. (2014). A mesoscale connectome of the mouse brain. Nature, 508(7495), 207–214. 10.1038/nature13186

32. Oscarsson, O. (1979). Functional units of the cerebellum - sagittal zones and microzones. Trends in Neurosciences, 2, 143–145.

33. Ozden, I., Sullivan, M. R., Lee, H. M., & Wang, S. S. H. (2009). Reliable coding emerges from coactivation of climbing fibers in microbands of cerebellar Purkinje neurons. Journal of Neuroscience, 29(34), 10463–10473. 10.1523/JNEUROSCI.0967-09.2009

34. Pisano, T. J., Dhanerawala, Z. M., Kislin, M., Bakshinskaya, D., Engel, E. A., Hansen, E. J., Hoag, A. T., Lee, J., de Oude, N. L., Venkataraju, K. U., Verpeut, J. L., Hoebeek, F. E., Richardson, B. D., Boele, H. J., & Wang, S. S. (2021). Homologous organization of cerebellar pathways to sensory, motor, and associative forebrain. Cell Rep, 36(12), 109721. 10.1016/j.celrep.2021.109721

35. Proville, R. D., Spolidoro, M., Guyon, N., Dugue, G. P., Selimi, F., Isope, P., Popa, D., & Lena, C. (2014). Cerebellum involvement in cortical sensorimotor circuits for the control of voluntary movements. Nat Neurosci, 17(9), 1233–1239. 10.1038/nn.3773

36. Ragan, T., Kadiri, L. R., Venkataraju, K. U., Bahlmann, K., Sutin, J., Taranda, J., Arganda-Carreras, I., Kim, Y., Seung, H. S., & Osten, P. (2012). Serial two-photon tomography for automated ex vivo mouse brain imaging. Nature Methods, 9(3), 255–258. 10.1038/nmeth.1854

37. Raymond, J. L., & Medina, J. F. (2018). Computational Principles of Supervised Learning in the Cerebellum. Annu Rev Neurosci, 41, 233–253. 10.1146/annurev-neuro-080317-061948

38. Rossi, L. F., Harris, K. D., & Carandini, M. (2020). Spatial connectivity matches direction selectivity in visual cortex. Nature, 588(7839), 648–652. 10.1038/s41586-020-2894-4

39. Sathyamurthy, A., Barik, A., Dobrott, C. I., Matson, K. J. E., Stoica, S., Pursley, R., Chesler, A. T., & Levine, A. J. (2020). Cerebellospinal Neurons Regulate Motor Performance and Motor Learning. Cell Rep, 31(6), 107595. 10.1016/j.celrep.2020.107595

40. Schultz, S. R., Kitamura, K., Post-Uiterweer, A., Krupic, J., & Häusser, M. (2009). Spatial pattern coding of sensory information by climbing fiber-evoked calcium signals in networks of neighboring cerebellar Purkinje cells. Journal of Neuroscience, 29(25), 8005–8015. 10.1523/JNEUROSCI.4918-08.2009

41. Schwarz, L. A., Miyamichi, K., Gao, X. J., Beier, K. T., Weissbourd, B., DeLoach, K. E., Ren, J., Ibanes, S., Malenka, R. C., Kremer, E. J., & Luo, L. (2015). Viral-genetic tracing of the input-output organization of a central noradrenaline circuit. Nature, 524(7563), 88–92. 10.1038/nature14600

42. Tervo, D. G. R., Hwang, B. Y., Viswanathan, S., Gaj, T., Lavzin, M., Ritola, K. D., Lindo, S., Michael, S., Kuleshova, E., Ojala, D., Huang, C.-C., Gerfen, C. R., Schiller, J., Dudman, J. T., Hantman, A. W., Looger, L. L., Scharf, R., & Kim, D. S. (2016). A designer AAV variant permits efficient retrograde access to projection neurons. Neuron, 92(2), 372–382. 10.1016/j.neuron.2016.09.021

43. Thanawalla, A. R., Wilcox, O., Rhee, E., Jiang, J., Huang, K. W., Yusufi, R., Saklaway, D., Nagamori, A., Conner, J. M., Chen, A. I., & Azim, E. (2025). Cerebellar outputs for rapid directional refinement of forelimb movement. bioRxiv. 10.1101/2025.10.01.679895

44. Tran-Van-Minh, A., Ye, Z., & Rancz, E. (2023). Quantitative analysis of rabies virus-based synaptic connectivity tracing. PLoS One, 18(4), e0278053. 10.1371/journal.pone.0278053

45. Tyson, A. L., Rousseau, C. V., Niedworok, C. J., Keshavarzi, S., Tsitoura, C., Cossell, L., Strom, M., & Margrie, T. W. (2021). A deep learning algorithm for 3D cell detection in whole mouse brain image datasets. PLoS Comput Biol, 17(5), e1009074. 10.1371/journal.pcbi.1009074

46. Wagner, M. J., Kim, T. H., Kadmon, J., Nguyen, N. D., Ganguli, S., Schnitzer, M. J., & Luo, L. (2019). Shared Cortex-Cerebellum Dynamics in the Execution and Learning of a Motor Task. Cell, 177(3), 669–682 e624. 10.1016/j.cell.2019.02.019

47. Wagner, M. J., Savall, J., Hernandez, O., Mel, G., Inan, H., Rumyantsev, O., Lecoq, J., Kim, T. H., Li, J. Z., Ramakrishnan, C., Deisseroth, K., Luo, L., Ganguli, S., & Schnitzer, M. J. (2021). A neural circuit state change underlying skilled movements. Cell, 184(14), 3731–3747.e3721.

48. Wang, X., Novello, M., Gao, Z., Ruigrok, T. J. H., & De Zeeuw, C. I. (2022). Input and output organization of the mesodiencephalic junction for cerebro-cerebellar communication. J Neurosci Res, 100(2), 620–637. 10.1002/jnr.24993

49. Wickersham, I. R., Finke, S., Conzelmann, K. K., & Callaway, E. M. (2007). Retrograde neuronal tracing with a deletion-mutant rabies virus. Nat Methods, 4(1), 47–49. 10.1038/nmeth999

50. Winship, I. R., Pakan, J. M., Todd, K. G., & Wong-Wylie, D. R. (2006). A comparison of ventral tegmental neurons projecting to optic flow regions of the inferior olive vs. the hippocampal formation. Neuroscience, 141(1), 463–473. 10.1016/j.neuroscience.2006.03.057

51. Wolpert, D. M., Miall, R. C., & Kawato, M. (1998). Internal models in the cerebellum. Trends Cogn Sci, 2(9), 338–347. 10.1016/s1364-6613(98)01221-2

52. Zhu, J., Hasanbegovic, H., Liu, L. D., Gao, Z., & Li, N. (2023). Activity map of a cortico-cerebellar loop underlying motor planning. Nature Neuroscience. 10.1038/s41593-023-01453-x

53. Zingg, B., Chou, X. L., Zhang, Z. G., Mesik, L., Li, H., Keene, C. D., Zaghloul, K. A., Tao, H. W., & Zhang, L. I. (2017). AAV-mediated anterograde transsynaptic tagging: Mapping corticocollicular input-defined neural pathways for defense behaviors. Neuron, 93(1), 33–47.

54. Zingg, B., Peng, B., Huang, J., Tao, H. W., & Zhang, L. I. (2020). Synaptic Specificity and Application of Anterograde Transsynaptic AAV for Probing Neural Circuitry. J Neurosci, 40(16), 3250–3267. 10.1523/JNEUROSCI.2158-19.2020

